# Variability of the innate immune response is globally constrained by transcriptional bursting

**DOI:** 10.1101/2023.02.20.529223

**Authors:** Nissrin Alachkar, Dale Norton, Zsofia Wolkensdorfer, Mark Muldoon, Pawel Paszek

## Abstract

Transcription of almost all mammalian genes occurs in stochastic bursts, however the fundamental control mechanisms that allow appropriate single-cell responses remain unresolved. Here we utilise single cell genomics data and stochastic models of transcription to perform global analysis of the toll-like receptor (TLR)-induced gene expression variability. Based on analysis of more than 2000 TLR-response genes across multiple experimental conditions we demonstrate that the single-cell, gene-by-gene expression variability can be empirically described by a linear function of the population mean. We show that response heterogeneity of individual genes can be characterised by the slope of the mean-variance line, which captures how cells respond to stimulus and provides insight into evolutionary differences between species. We further demonstrate that linear relationships theoretically determine the underlying transcriptional bursting kinetics, revealing different regulatory modes of TLR response heterogeneity. Stochastic modelling of temporal scRNA-seq count distributions demonstrates that increased response variability is associated with larger and more frequent transcriptional bursts, which emerge via increased complexity of transcriptional regulatory networks between genes and different species. Overall, we provide a methodology relying on inference of empirical mean-variance relationships from single cell data and new insights into control of innate immune response variability.

## Introduction

Transcription of almost all mammalian genes occurs in bursts, during brief and random periods of gene activity. The patterns of temporal mRNA production in a single cell, and the overall mRNA (and protein) distribution in cellular populations, are controlled by transcriptional bursting, namely via the modulation of *burst size* and *burst frequency* [1–3]. The innate immune responses exhibit extreme variability at the single cell level, in comparison to other tissue systems [4–6], where only subsets of cells produce specific effector molecules, and thus are able to restrict pathogen growth [7]. This apparent level of variability poses a fundamental systems biology question; how do robust immune responses emerge from this heterogeneous transcriptional bursting process?

Recent advances have demonstrated key insights into regulation of transcriptional bursting. In general, the bursting kinetics are gene-specific and subject to regulatory control via cellular signalling events [3, 8–11] as well as genome architecture and promoter sequences [4, 12–16]. For example, core promoters control burst sizes, while enhancer elements modulate burst frequency to define cell-type specific [17] or circadian gene expression outputs [18]. Coordinated gene activity has also been shown to regulate mRNA outputs as a function of spatial position during development [19–21] as well as temporal immune responses [22]. The resulting cell-to-cell variability is a consequence of the stochastic processes governing signalling and transcription [23], but also reflects extrinsic differences between individual cells [24–27] or variability of the pathogen in the context of the innate immune response [7]. With individual genes exhibiting different levels of stimuli-induced heterogeneity, we are still lacking general understanding of how transcription is regulated at the single cell level.

Toll-like (TLR) receptor signalling constitutes one of the fundamental, evolutionarily conserved innate immune defence mechanisms against foreign threats [28, 29], yet exhibits substantial cell-to-cell variability [4–6, 30, 31]. We recently demonstrated that this overall TLR response to stimulation (or in general perturbation) is constrained through gene-specific transcriptional bursting kinetics [32]. By utilising single molecule Fluorescent in situ Hybridisation (smFISH), we established that the overall mRNA variability is linearly constrained by the mean mRNA response across a range of related stimuli. Variance (and in fact higher moments) of the mRNA distributions have been also shown to be constrained by the mean response in the developing embryo [19]. These analyses suggest that complex transcriptional regulation at a single cell level may be globally characterised by mean-variance relationships of gene-specific mRNA outputs, providing new ways to characterise response variability. While quantitative smFISH provides important insights, this approach is often limited by the number of genes, which can be investigated [8, 9, 32–37], therefore further analyses of global gene expression patterns [15, 17] are required to fully understand the underlying regulatory constraints.

Here we utilise scRNA-seq data on innate immune phagocytes stimulated with common TLR ligands, lipopolysaccharides (LPS) of Gram-negative bacteria upstream of TLR4 and viral-like double-stranded RNA (PIC) for TLR3 [4] to investigate the control of single cell gene expression heterogeneity of the innate immune responses. We analyse 2,338 TLR-response genes and demonstrate that they globally follow empirical linear mean-variance relationships, exhibiting a genome-wide spectrum of response variability levels characterised by the slope of the relationship. We show that linear relationships define different modes of individual gene response modulation with majority of the genes undergoing frequency modulation to TLR stimulation. Mathematical modelling of scRNA-seq count distributions using dynamic stochastic telegraph models of transcription of varied complexity levels, demonstrates that increased response variability is associated with larger and more frequent transcriptional bursts, which emerge via increased regulatory complexity. Finally, we show that linear mean-variance relationships capture evolutionarily differences in response variability across pig, rabbit, rat, and mouse and predict transcriptional bursting modulation between species. Overall, our data demonstrate the utility of empirical mean-variance relationships in providing new insights into control of transcriptional variability in the innate immune response.

## Results

### TLR-induced mRNA responses exhibit linear mean-variance trends

To globally investigate the control of transcriptional bursting in the TLR system relationships we used existing scRNA-seq data from mouse phagocytes either untreated or stimulated with LPS and PIC for 2, 4 and 6 hours [4]. The dataset contains unique molecular identifier (UMI) mRNA counts for 53,086 cells and 16,798 genes across 20 experimental conditions including replicates, of which 2,338 genes were identified as TLR-dependent (see Fig. 1A for correlation of sample mean and variance across all datasets, and Materials and Methods for data processing). We previously showed that the gene-specific variability can be defined by the slope of the mean-variance relationship [32]. To test this phenomenon globally, for each of the 2,338 TLR-inducible genes, the sample mean (μ) and variance (σ^2^) relationship was fitted using robust linear regression (*σ*^2^ = *αμ* + *α*_0_), yielding 2,133 genes with a significant regression slope (*p*-value < 0.05, Fig. 1B). Of those, 1,551 (66% of all TLR-inducible genes) genes, referred here as high confidence genes, were characterised by coefficient of determination *R*^2^ > 0.6 (Fig. 1C, see also Table S1 for list of genes and fitted relationships). Overall, the distribution of fitted slopes across the high confidence genes varied over 3 orders of magnitude, with 1,067 genes (69% of high confidence genes) characterised by slope α>1 and 627 (40%) α>3, indicative of predominant non-Poissonian transcription (where one would expect α=1 and *α*_0_ = 0) (Fig. 1D). 61 genes (4%) were characterised by α> 5 and 28 (2%) by a α>10, highlighting genes with the highest level of expression variability (across a range of TRL responses, Fig. S1A). Among the high variability genes (α>5) we found C-C motif chemokine ligands (*Ccl*) *2, 3, 4, 5, 17*; C-X-C motif ligands (*Cxcl*) *9* and *10*, as well as cytokines including Interleukin 1 α (*IL1a*), *IL1b*, *IL10*, *IL12b* and Tumour Necrosis Factor α (*Tnfa*) (see Fig. 1E for individual gene fits). The most variable gene in the dataset was the immunoglobulin subunit *Jchain* with α=1372 (Fig. S1D), substantially more than the 2^nd^ most variable *Ccl5* (α=72). While the range of the mRNA output among high confidence genes varies over 3 orders of magnitude (Fig. S1B), we found that LPS induced more robust activation than PIC in terms of average expression (Fig. 1F). Overall, this analysis demonstrates that TLR-induced mRNA responses globally exhibit empirical linear mean-variance relationships.

**Figure 1.**
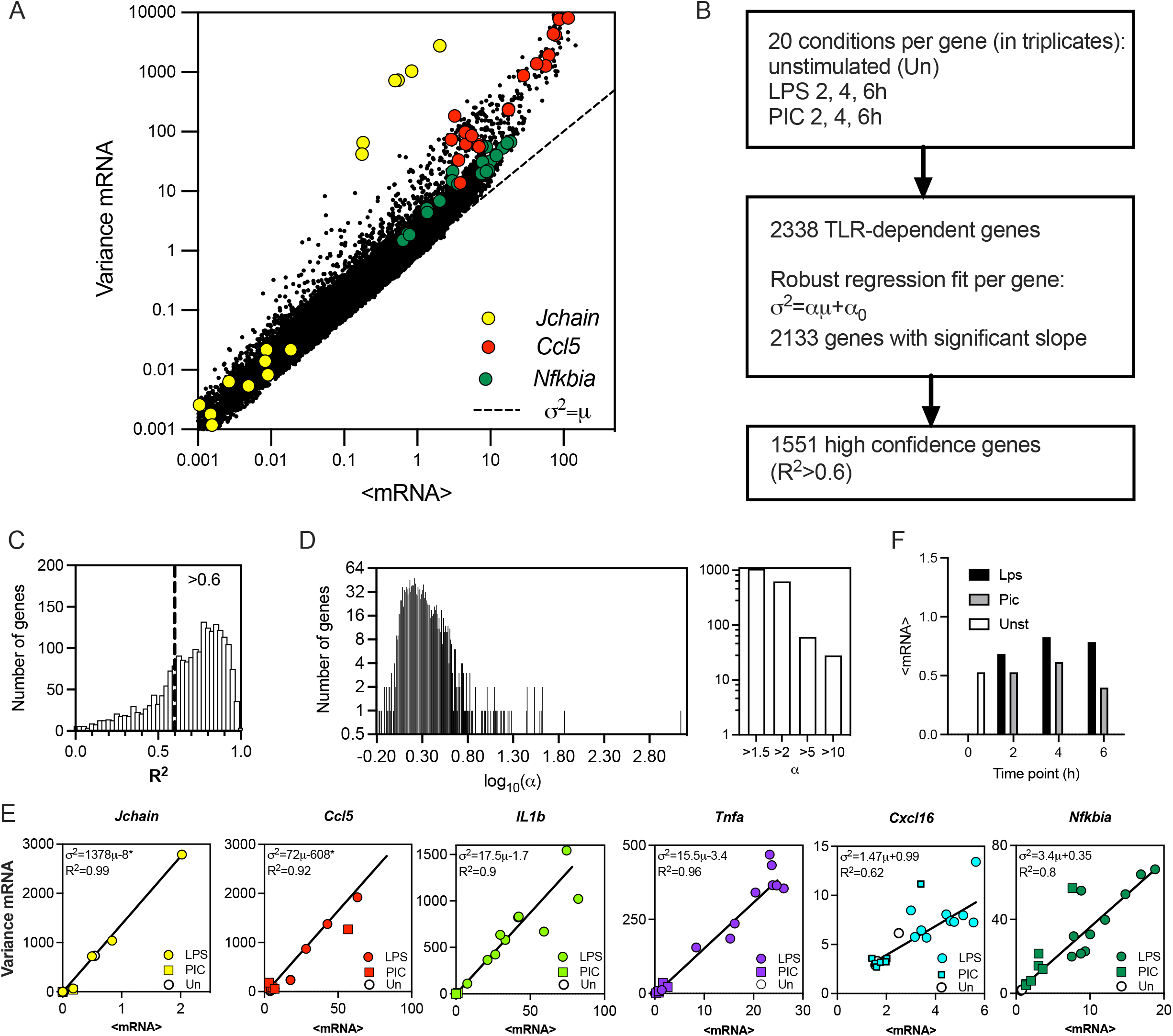
TLR-induced transcriptional variability is linearly constrained. **A**. Overall variability in the scRNA-seq dataset [4]. Shown is the scatter plot of the sample mean (μ) and variance (σ^2^) calculated for 2340 TLR-dependent genes across 20 experimental conditions. Data points corresponding to *Jchain*, Ccl5 and *Nfkbia* highlighted in yellow, red, and green, respectively. Broken line indicates *μ=σ^2^* line. **B**. Schematic description of the fitting protocol. **C**. Histogram of coefficient of determination (*R^2^*) for 2,133 gene fits characterised by a significant regression slope (*p*-value < 0.05). *R^2^* = 0.6 broken line corresponds to the high confidence gene cut-off. **D**. Distribution of the fitted regression slopes for the 1,551 high confidence gene set. Histogram of the fitted slopes shown on the left. Number of genes with different slope range shown on the right. **E**. Fitted mean-variance relationships for a subset of genes. Shown are the individual datapoints (LPS, PIC and unstimulated) as well as fitted regression line with a fitted equation (* denotes statistically significant intercept, *p*-value < 0.05) and the coefficient of determination (*R^2^*). **F**. Mean mRNA counts across treatments (LPS, PIC) and time (0, 2, 4, 6 h) for the 1,551 high confidence genes.

### Patterns of transcriptional bursting modulation underlie TLR response heterogeneity

Having established the linear relationships relating the gene-specific transcriptional variability to mean expression, we sought to study global properties of transcriptional bursting underlying these trends. We used moment estimators of the underlying scRNA-seq count distributions to calculate bursting characteristics, such that burst size *b_s_=σ*^2^/*μ* (i.e., the Fano factor) and burst frequency *b_f_=μ/b_s_*, which measure the departures from Poissonian mRNA production [1, 3, 11, 18]. Given the empirical linear constraint, *σ*^2^ = *αμ* + *α*_0_, the burst size and burst frequency become analytical functions of the mean mRNA expression such that *b_s_=α_0_/μ* + *α* and *b_f_=μ^2^/(α_0_+ aμ)* (Fig. 2A). In a special case when *α_0_=0*, burst size is constant (independent of the mean expression μ) and equal to the slope of the mean-variance line α, while the frequency increases linearly with *μ* and is proportional to *1/α* [32]. However, the overall behaviour does depend on the intercept (see Fig. S1C for sensitivity analyses); for α_0_>0, the burst frequency converges monotonically to μ/α (i.e., the limiting case for *α_0_=0*), while the burst size converges to α (from ∞ at *μ=*0) as the mean expression *μ* increases (Fig. 2A in blue). For α_0_<0 (Fig. 2A, in red), the relationship can only be defined for *μ>/α_0_/^/^α* such that burst size increases monotonically (and converges to α), while the burst frequency has a local minimum for μ*=*2/α_0_/^/^α* equal to *4/α_0_/^/^α^2^*, eventually converging to the limiting case *μ/α*.

**Figure 2.**
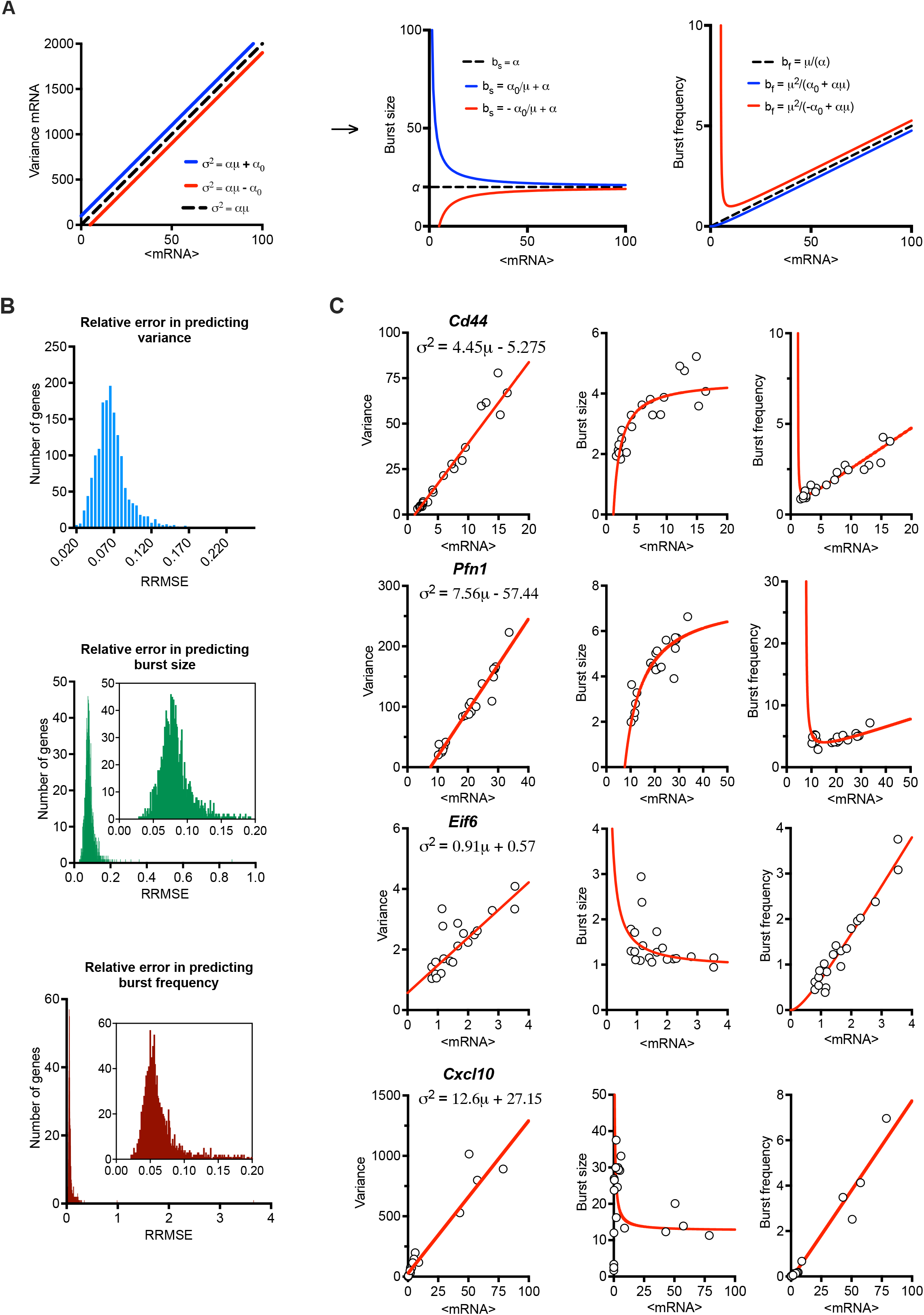
Mean-variance relationships constrain transcriptional bursting characteristics. **A**. Theoretical burst size and frequency characteristics. (Left) Simulated mean variance relationships with positive (in blue, α=20, α_0_=100) and negative (in red, α=20, α_0_= −100) intercepts, respectively. (Middle & Right) Derived burst size and frequency modulation schemes for corresponding parameter values calculated using moment estimators. A special case of α=20, α_0_=0 is shown in broken line. **B**. Global modulation of transcriptional busting. Shown is the comparison between fitted mean-variance relationship and derived theoretical burst size and frequency modulation schemes vs. experimental data. Shown is distribution of relative root mean square error (RRMSE) of 1,551 high confidence genes. **C**. Modulation schemes for *Cd44, Pfn1, Eif6* and *Cxcl10* genes. Shown is the comparison between theoretical relationships based on fitted mean-variance relationships (in red) and corresponding estimates from data (open circles). Equations for fitted mean-variance relationships highlighted in the top left panel, respectively.

We calculated the theoretical bursting modulation schemes for the 1,551 high confidence genes and compared these to the moment estimators of the burst size and frequency from the data (Fig. 2B). We found that the average relative root mean square error (RRMSE, see methods) of the mean-variance fit in relation to data was 0.07±0.02%, where 1,431 genes had an error smaller than 0.1%. In comparison, the average error for the burst size modulation was 0.08±0.03% (with 1281 genes having an error smaller than 0.1%), while the average error for the burst frequency modulation was 0.07±0.1% (with 1,389 genes having an error smaller than 0.1%). Given their empirical nature, the predicted theoretical trends are in good agreement with the changes of burst size and frequency observed in the data. Profilin 1 (*Pnf1)* and *Cd44* are example genes characterised by intercept α_0_<0, while the genes encoding eukaryotic translation initiation factor 6 (*Eif6*) and *Cxcl10* had α_0_>0 (Fig. 2C). *Jchain* is an example of a gene with a good mean-variance fit, but one of the poorest fit in terms of bursting frequency, which might be due to limited sample size and its profound variability (Fig. S1D). Of the 1,551 high confidence genes, 430 genes had a significant intercept (*p*-value < 0.05) in the regression fit, with 414 characterised by negative and 16 positive intercepts (Fig. S1E). These in part reflect the empirical nature of these trends and the limited sample size, especially for those genes where *α_0_* is small (in relation to variance), for example *Cxcl10* (Fig. 2C). However, many genes, including *Pnf1* and *Eif6* exhibit substantial basal expression in untreated cells [8], resulting in either elevated or reduced variability (in relation to true zero) as captured via non-zero intercept in the regression fit [32].

### Gene specific bursting exhibits different modes of response modulation

The linear mean-variance relationships reflect the constrained changes of burst size and burst frequency required to regulate response variability as shown in their derived analytical functions of the mean mRNA expression. To understand the modulation of transcriptional bursting, we first calculated fold changes of burst size vs. burst frequency across the range of mean expression calculated for individual response genes (Fig. 3A). We found that 1,015 out of the 1,551 high confidence genes exhibit 2 times more fold changes in burst frequency than burst size. This suggests a predominant frequency modulation, in agreement with recent analyses of LPS-induced macrophages [22]. However, we also found 48 genes exhibiting fold changes in burst size 2 times more than burst frequency, while 389 exhibited comparable modulation of both burst size and burst frequency. To study the transcriptional bursting modulation more systematically, we derived an analytical relationship between the burst size and frequency (independent of the mean mRNA expression) based on the linear constrains (Fig. 3B). The general relationship is given by *b_f_=α_0_/(b_s_(b_s_-α))*, where α_0_ can take positive or negative values. When α_0_>0, we have an inverse relationship between the burst size and frequency, which asymptotically approaches zero, as the burst size approaches infinity. It is also worth mentioning that, in this case, the function is undefined for values of burst size smaller than or equal to *α* (Fig. 3B, in blue), reflecting a biological limit of burst size and frequency for genes following this modulation trend. We found that 315 genes (out of the 1,551 high confidence genes) exhibited such an inverse relationship, with all genes exhibiting higher frequency than burst size modulation (see Fig 3C for specific genes and Fig 3D and Table S2 for global analysis). For the case when α_0_<0, linear constrains define a non-monotonic relationship between the burst size and frequency on the interval (0,α) with a local minimum at 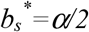, and frequency diverging to infinity as burst size tends towards α or is close to 0 (Fig. 3B, in red). From the case α_0_<0, three patterns of bursting modulation can be distinguished; the burst frequency and size exhibit either inverse relationship, where the frequency increases and burst size decreases (for 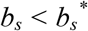) or concurrent increases 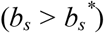. In addition, we define a U-shape relationship such that 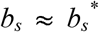 where both inverse and concurrent relationships are possible (i.e., 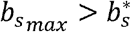 and 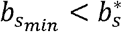 per gene). We found that out of the 1236 genes characterised by α_0_<0, most genes (999) exhibited predominant frequency modulation following either a U-shape or a concurrent relationship, while 237 genes showed higher burst size modulation and was mostly associated with U-shape trends (Fig. 3C and D). It is worth mentioning that all 7 genes confirming an inverse trend showed predominant burst size modulation. Overall, these analyses demonstrate different modes of the transcriptional bursting modulation of TLR-stimulated genes, albeit with predominant regulation via burst frequency.

**Figure 3.**
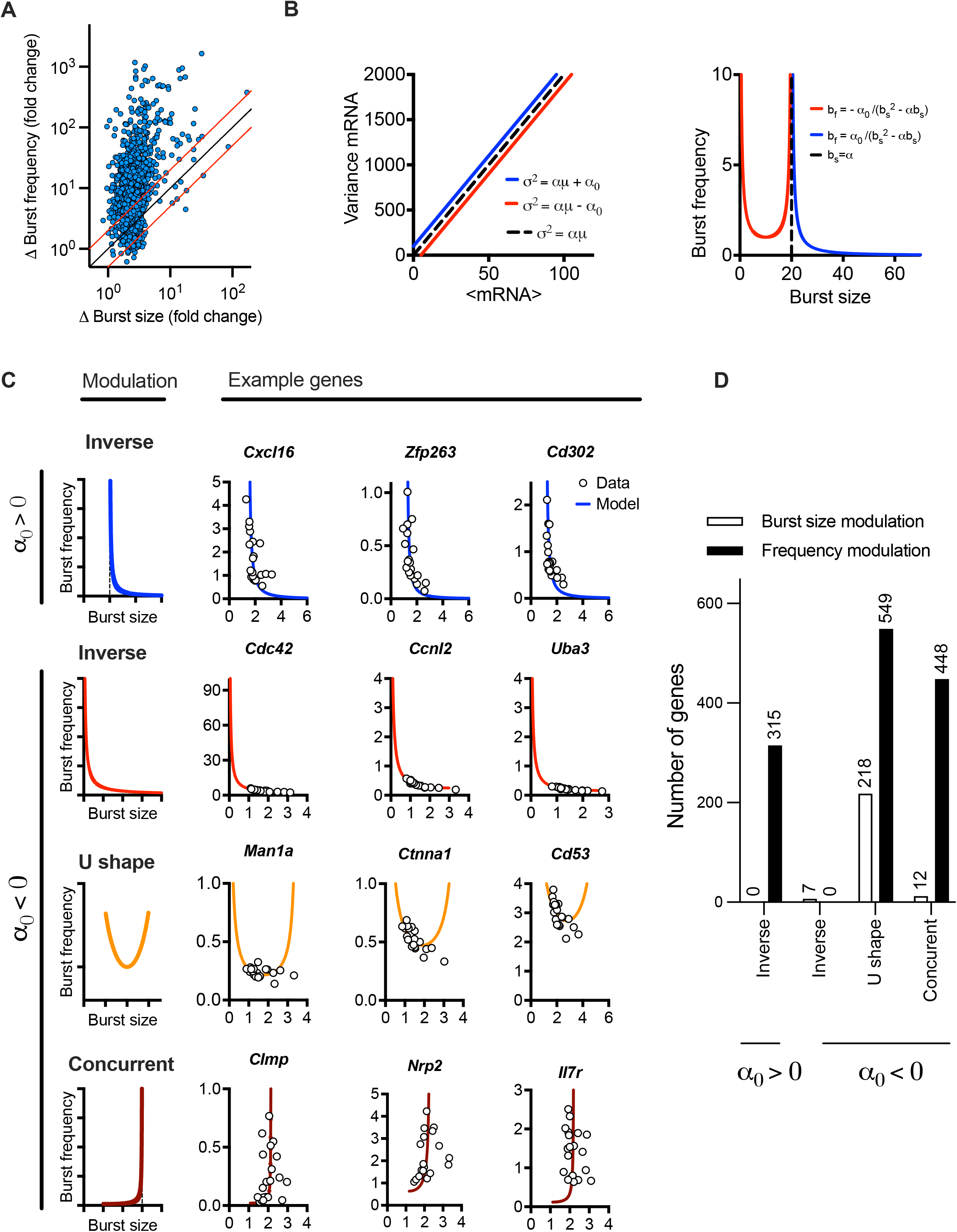
LPS-induced gene expression undergoes different modes of transcriptional bursting. **A**. Relative changes of burst size and burst frequency. Shown is the relative fold change of burst size and frequency calculated across the individual range of mean expression for 1,551 high confidence genes (in blue circles). Identity line depicted in black, two-fold change highlighted in red. **B**. Theoretical relationship between burst size and frequency. (Left) Simulated mean variance relationships with positive (in blue, α = 20, α_0_ = 100) and negative (in red, α = 20, α_0_ = −100) intercepts, respectively. (Right) Burst size and frequency modulation schemes for corresponding parameter values calculated using moment estimators. A special case of α = 20, α_0_= 0 shown in broken line. **C**. Modulation of burst size and frequency across a range of individual genes. Shown are inverse relationship (*α_0_>0*) in blue as well as inverse, U-shape and concurrent relationships (*α_0_<0*). Relationship predicted from linear constraints in solid lines and corresponding estimates from experimental data in open circles. U-shape numerically defined as maximum burst size value > *α/2* and minimum burst size value < α/2 across conditions. **D**. Prevalence of different modulation schemes across 1,551 high confidence genes. Definition of the mode as in C, dominant modulation defined by absolute difference in the burst size vs. frequency changes across the respective range of mean expression (as in A).

### Increased response variability is associated with complex transcriptional regulation

The distribution of fitted regression slopes varying over 3 orders of magnitude demonstrate a wide range of response variability among individual TLR-induced genes (Fig. 1D). While we have demonstrated that individual genes exhibit different modes of transcriptional bursting characteristics to regulate responses to stimulation, we wanted to understand the control of variability in the system more mechanistically. A well-established mathematical description of mRNA production involves a 2-state telegraph model (Fig. 4A), where gene activity changes randomly between “off’ and “on” states, with mRNA transcription occurring in the “on” state [1, 3, 18, 36]. The associated parameters are gene activity rates (*k_on_* and *k_off_*) as well as rate of mRNA transcription (*k_t_*) and degradation (*k_d_*) (Nicolas et al., 2018). Although the 2-state telegraph model has been widely used in the past to model mRNA count data, more complex structures are often required to capture additional complexity associated with multiple regulatory steps, combinatorial promoter cycling and transcriptional initiation [12, 38]. We previously showed that heterogenous *Il1β* mRNA transcription requires more regulatory steps than that of *Tnfα* [32]. We therefore hypothesised that TLR response variability is linked with the complexity of the transcriptional regulation. To test this hypothesis, we introduced a 3-state stochastic model, which assumes sequential promoter activation between “off”, “intermediate” and “on” states, equivalent to promoter cycling [12, 38], with transcription occurring in the “intermediate” (*I*) state as well as in the “on” state, characterised by 5 transition rates (*t_on_, t_off_, k_on_, k_o_ff*and *k_c_*), 2 transcription rates (*k_ti_* and *k_t_*), and a degradation rate *k_d_* (Fig. 4A).

**Figure 4.**
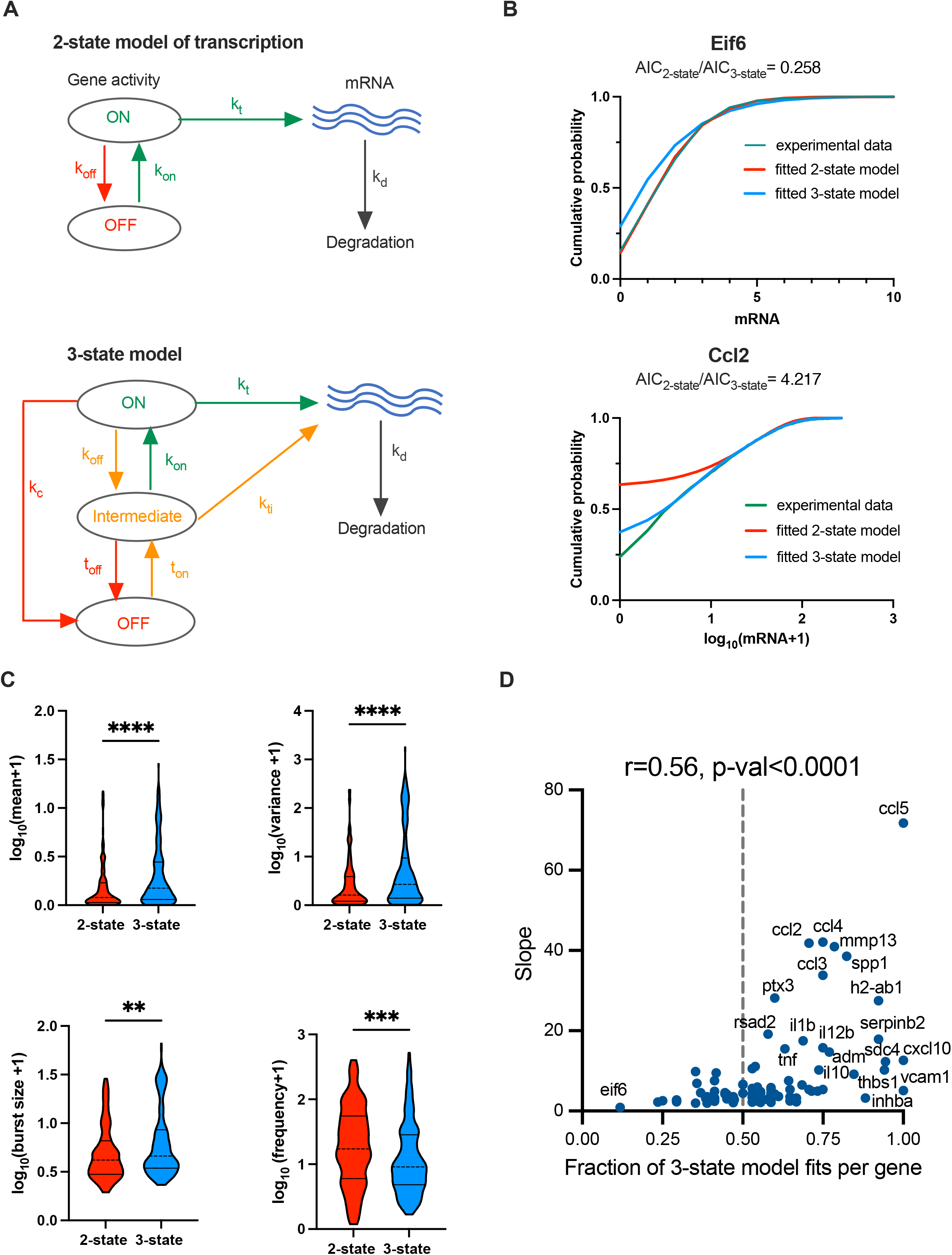
TLR response variability is associated with regulatory complexity. **A**. Schematic representation of the 2-state and 3-state models of transcription. **B**. Comparison between the fitted and measured mRNA counts distributions. Shown are cumulative probability distribution of data (in green) vs. the corresponding 2-state and 3-state stochastic model fits (in red and blue, respectively) for representative condition for *Eif6* (PIC, 4h, replicate 3) and *Ccl2* (LPS, 2h, replicate 2) genes. **C**. Analysis of transcriptional bursting across high coverage genes and conditions fitted by 2-state vs 3-state models. Shown is the comparison between best fit 2-state and 3-state models in terms of mean mRNA expression, variance, burst size and frequency from experimental data. Best fit defined by *AIC_best model_<0.5AIC_2nd best_* (from Fig. S3B). Burst size and frequency calculated per condition using moment estimators. Statistical significance assessed with Mann-Whitney test (** *p*-value<0.01, **** *p*-value <0.0001). **D**. Relationship between slope of the mean-variance relationship and fraction of 3-state model fits for high coverage genes. Fraction of 3-state model fits per gene defined by the number of conditions with *AIC_3-state model_<AIC_2-state_* over all conditions per gene. Broken line indicates 0.5, *r* denotes Spearman correlation coefficient.

We first used a profile likelihood approach [17, 39] to fit the measured scRNA-seq count distributions assuming steady state kinetics of the 2-state model (the so called Beta-Poisson model) for the 1,551 high confidence genes, each across 20 treatment datasets (Table S3). Values of kinetic parameters were inferred for 7,804 of 31,020 datasets (~25% across 1519 genes), which in general corresponded to genes characterised by larger expression, in comparison to those that failed to fit (Fig. S2A). The fitted parameter values (*k_on_, k_off_* and *k_t_*, expressed in units per degradation half-life) varied over 3 orders of magnitude across all genes and datasets (Fig. S2B). In general, gene inactivation rates (*k_off_*) were greater than activation rates (*k_on_*) (Fig. S2C), consistent with intermittent transcriptional kinetics [3, 13, 17]. While the Beta-Poisson model explicitly assumes a steady-state (and does not make any assumptions about mRNA half-life), we wanted to account for the underlying dynamical stochastic processes and corresponding temporal mRNA production and decay [34]. However, it was not computationally feasible to fit all genes across all scRNA-seq datasets, we therefore identified on a subset of 99 high confidence genes for which at least 10 datasets were fitted using a Beta-Poisson model (Fig. S2D). Of these, 96 had an existing measurement of mRNA half-life (which is required for dynamical model fitting) in LPS-stimulated bone marrow derived macrophages [40, 41] or other cell models. The resulting 96 high coverage genes included 51 of 100 most variable genes (as defined by the fitted regression slope) and 60 of 100 most expressed genes including chemokine family *Ccl5, Ccl4, Ccl3, Ccl2* as well as *IL1b* and *TNFa* (Fig. S2D, E and F, see Table S3 for a list of genes, half-lives and fitted relationships).

We used a genetic algorithm to fit dynamical 2-state and 3-state stochastic models across 20 individual datasets (LPS and PIC stimulation at 0, 2, 4, 6 h time-course across replicates) for the 96 high coverage genes (see Material and Methods). We then applied the Akaike information criterion (AIC) [42] to select models that accurately fitted the measured mRNA distributions and compare the quality of the three models per condition in order to determine the best-fit model, noting that the lower AIC value corresponds to the better model fit. In general, we found that Beta-Poisson model, the least constrained model, fitted better than dynamical models (805 out of 1210 conditions (i.e., treatment and replicates) favoured Beta-Poisson model based on their AIC values, Fig. S3A and B). The more constrained dynamical 2-state model provided a best fit for 170 conditions, while the 3-state model best captured 235 conditions (and 30 and 57, respectively when using a more stringent criterion of two-fold AIC change, Fig. S3B). When comparing 2-state with 3-state model directly and assuming a two-fold AIC change between the two models, there were 141 out of 1507 conditions that favoured the 2-state model, while the opposite was true for 266 conditions (see Fig. S3C for other thresholds). For example, 2-state model recapitulated PIC-treated *Eif6* mRNA count distribution (at 4 h) better than a 3-state model, as reflected by the *AIC_2-state_<AIC_3-state_*. In turn, the 3-state model better recapitulated the LPS-treated *Ccl2* distribution (at 2 h) spanning almost over 3 orders of magnitudes (Fig. 4B). The number of 2-state-and 3-state model fits was not strongly related to the treatment, time point or in fact biological replicates, although LPS had 155 conditions more fitted with 3-state than 2-state model (Fig. S3D).

The 141 2-state model fits were characterised by *k_on_=0.02 ±0.01* min^-1^ (half-time of 35 mins) on average, and off rates averaging *k_off_=0.74±0.25* min^-1^ (half-time of 1 min), with average transcription rate *k_t_=1.23± 4.44* mRNA min^-1^, indicative of ‘bursty’ kinetics (Fig. S4A). The ‘*on*’ rate showed significant positive correlation with the variance of the corresponding count distributions (*r=0.48*), demonstrating that a faster ‘*on*’ switch contributes towards increased response variability. The 266 3-state model fits were also characterised by relatively slow average ‘*on*’ rates (*t_on_=0.036 ±0.13* min^-1^ and *k_on_=0.33 ±0.32* min^-1^) in relation to the ‘*off*’ rates (*t_off_=0.74± 0.26* min^-1^, *k_off_=0.44± 0.36* min^-1^ and *k_c_=0.50± 0.36* min^-1^, Fig. S4B). The mRNA count variance was correlated positively with *t_on_* rate (i.e., transition to intermediate state, *r=0.33*) as well as with transcription rates in ‘on’ and ‘intermediate’ states (r>0.4). In comparison to the 2-state model, the transcription rate in the ‘on’ state was significantly higher (*k_t_=7.63± 13.05* mRNA min^-1^) indicative of larger burst sizes (Fig. S4C and D).

We then asked if the level of variability is linked with the model complexity. We found that scRNA-seq count distributions fitted with the 3-state model were characterised by greater variability than those corresponding to the 2-state model (see Fig. 4C and Fig. S4D for less stringent model selection thresholds). In agreement, the 3-state-model fits were associated with significantly larger burst size and lower burst frequency than that of the 2-state model fits, consistent with more heterogenous bursting kinetics across the relevant conditions. Finally, we analysed model selection across individual high coverage genes rather than corresponding conditions; we found the fraction of conditions explained by one model changes between individual genes (e.g., 3-state model fitted 3 out of 20 for *Eif6*, 10 out of 20 for *Ccl5* and all conditions for *Vcam1* Fig. 4D). Our interpretation of this is that as the mRNA responses increase, a more complex regulatory structure is required to capture the underlying distribution. We found that, for the high coverage genes, the fraction of conditions explained by the 3-state model correlated well (*r=*0.56, *p*-value < 0.0001) with the slope of mean-variance relationship, and thus response heterogeneity (Fig. 4D). Overall, this demonstrates that while increased heterogeneity involves larger and infrequent bursts (in comparison to homogenous responses), this is underlined by increased complexity of the transcriptional regulatory network.

### Linear relationships capture evolutionary changes of response variability

Previous work highlighted the relationship between evolutionary response divergence of innate immune genes and their cell-to-cell variability, with highly divergent genes exhibiting more variability [4]. However, the changes in patterns of transcriptional bursting during evolution is still poorly understood. We proposed that by comparing the linear mean-variance relationships across species, the variations in transcriptional bursting patterns that develop through evolution could be better understood. Specifically, if the evolutionary changes in response variability can be captured by a fold-change *k* in the slope of the relationship, then the increased variability is predicted to be due to increased burst size and reduced burst frequency by a factor *k*, respectively (Fig. 5A).

**Figure 5.**
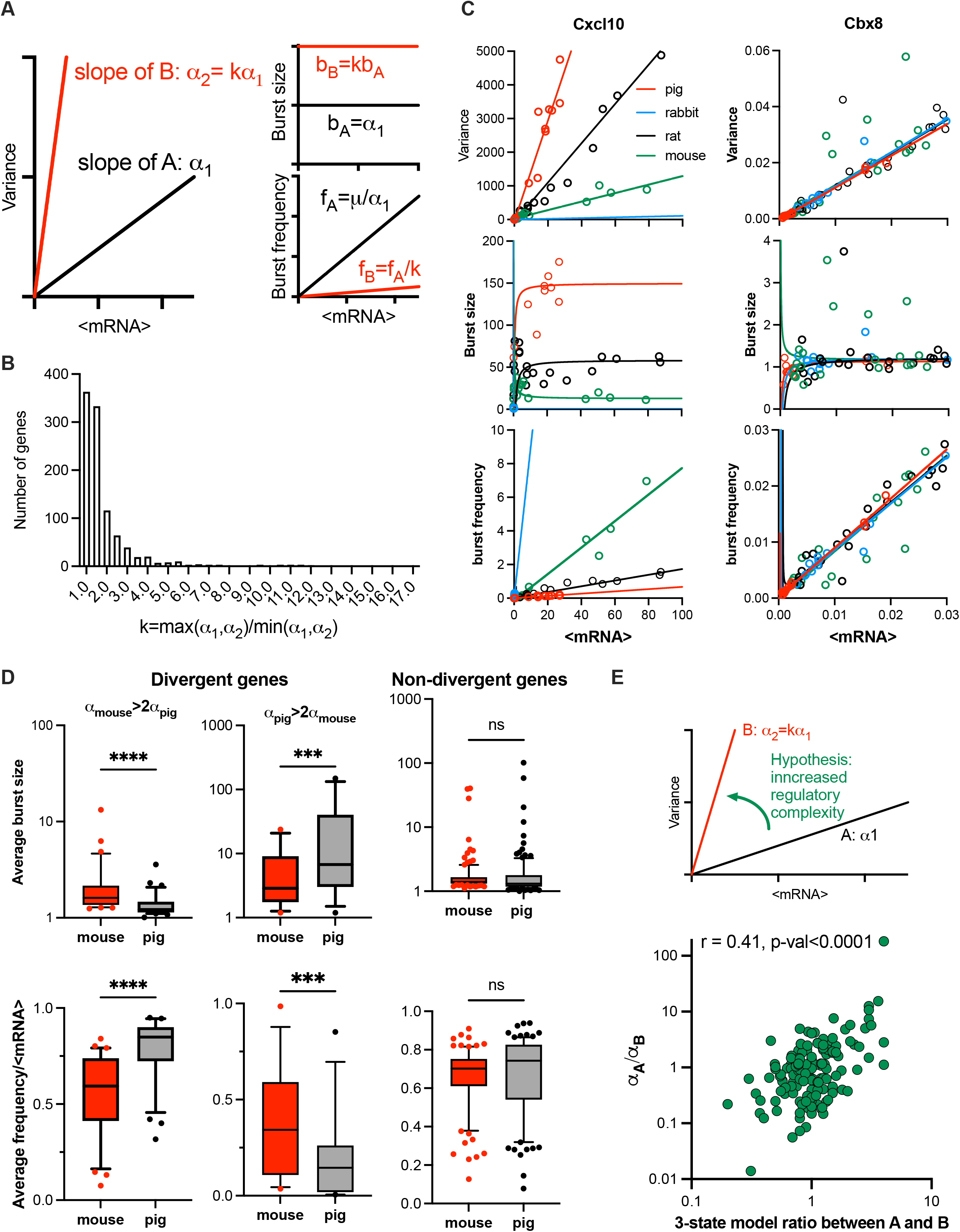
Evolutionary control of TLR response variability. **A**. Schematic representation of response variability during evolution for putative species A and B. Shown are mean variance relationships corresponding to slopes (*α_1_* and *α_2_=kα_1_*) and the predicted burst size (*b*) and frequency *(f)* modulation schemes for corresponding parameter values calculated using moment estimators. **B**. Histogram of the slope ratio *k* calculated for the 169 orthologue genes across all pairwise comparisons between mouse, rat, rabbit and pig. *k=max(α_1_,α_2_)/min(α_1_,α_2_)*, where α_1_ and α_2_ denote slopes of the fitted mean-variance relationships for each pair of species per gene. **C**. Modulation schemes for *Cxcl10* and *Cbx8* genes. Shown is the comparison between theoretical relationships based on the fitted mean-variance relationships (in solid lines, colour-coded by species) and corresponding moment estimates for burst size and frequency from experimental data (circles). **D.** Analysis of burst size and frequency for divergent and non-divergent mouse and pig TLR-response genes. Shown are box plots of average burst size and mean-normalized frequency per gene stratified into divergent (α_mouse_> 2α_pig_ or α_pig_>2α_pig_) and complementary non-divergent groups (31, 15 and 123 orthologue genes, respectively). Statistical significance assessed with a paired Wilcoxon test (**** *p*-value < 0.0001, *** *p*-value < 0.001, ns not significant). **E**. Change of variability between species is associated with regulatory complexity. Top: Schematic representation of the hypothesis. Bottom: Relationship between the slope ratio (α_A_/α_B_) estimated for 146 pairwise comparisons between 28 fitted orthologue genes for mouse, rat, rabbit and pig; and the corresponding ratio between species A and B of the number of conditions per gene with 3-state model fitting better than 2-state model. Absolute difference in AIC of the two models was used for model selection. Shown is the Spearman correlation coefficient *r* and a *p*-value for *r* > 0.

The relationship between the mean and variance of the single cell mRNA counts was studied in data for four mammalian species from Hagai *et al*. (2018): mouse, rat, pig, and rabbit, in cells either untreated or treated with LPS or PIC for 2, 4 and 6 h (see methods and Table S4 for species specific number of conditions per gene ranging from 12 to 21). We found that from the 2,338 LPS response genes, a subset of 218 genes with one-to-one orthologues showed response to treatment in all four species (Fig. S5A). 78% of fitted mean-variance relationships for the 218 genes were characterised by *R^2^* > 0.6, including 102 genes in all four species and 169 in at least three species. To characterise the divergence in response variability we performed species pairwise comparison between the fitted regression slopes of the 169 genes subset (Table S5). Out of this subset 21 genes including chemokines *Ccl2, Ccl4, Ccl5* and *Cxcl10* (Fig. 5B and Fig S5B), had all 6 possible pairwise comparisons showing significant differences, indicating divergence in TLR response variability between each of the two species. 5 significant FDR values (difference in three out of four species) were obtained for 49 genes including chemokines *Ccl20, Ccl3, MMP9* (Fig S5B) and cytokines *Il1a, Il10* and *Il27* indicating significant differences in response variability. On the other hand, no significant differences were obtained between any of the four slopes in 7 genes, including a transcriptional repressor Chromobox Protein Homologue 8 (*Cbx8*, Fig. 5B). In agreement, a distribution of slope ratios calculated across all pairs of species for the 169 genes (Fig. 5C and Table S6) revealed 49 pairs with *k > 5* and 258 pairs with *k > 2*, indicating substantial changes of the response variability between specie*s*, including the chemokine and cytokine genes. Conversely, 54% of slope ratios (549 out of total 1014 genes) were smaller than 1.5, indicative of conserved variability. The inflammatory chemokines were shown previously to rapidly evolve in mammals and other vertebrates with clear differences in expression between closely related species [43, 44]. Moreover, gene duplication of the CC chemokine ligands can result in different copy numbers of these genes between individuals [45], further increasing the divergence in expression. Importantly, our analyses specifically capture changes of response variability and suggest a statistical relationship of these changes with the generic evolutionary divergence (see Materials and Methods) of gene expression response (Fig. S5C).

To validate the predicted changes in transcriptional bursting during evolution (Fig. 5A), we first calculated the theoretical modulation schemes for all the 169 evolutionary genes across species and compared these to the moment estimators of the burst size and frequency from the data (Fig. S5D). We found that the average RRMSE of the mean-variance fit in relation to data was 0.06±0.05% across all species, where 90% genes had an error smaller than 0.1%. In comparison, the average error for the burst size predictions was 0.08±0.05%, while the average error for the burst frequency predictions was 0.05±0.04%. The predicted theoretical trends are in good agreement with the observed changes of burst size and frequency. For example, *Cxcl10* exhibits concurrent changes of the burst size and frequency, the level of which is determined by the slope of the relationships, while *Cbx8* exhibits the same modulation across species (Fig. 5C). In addition, our predictions of species-specific modulation scheme are based not only on the slope α, but also the mean-variance intercept, which we previously showed may affect the bursting relationships (Fig. 2A and Fig. S1C). We therefore investigated if the difference of the slopes alone is sufficient to predict modulation of bursting characteristics across species (Fig. 5A). We stratified the 169 orthologous genes into divergent and non-divergent subsets, with the divergence threshold defined by a 2-fold change in the slope of the mean-variance relationships. The divergent subset included 31 genes exhibiting higher slope in mouse, and 15 in pig (Fig. S5E). We found that divergent genes, associated with increased response variability, exhibited significantly higher average burst sizes (as calculated across all corresponding conditions) and reciprocally lower normalised burst frequency when compared between the two species (Fig. 5D). In contrast, the non-divergent genes showed no significant differences in the burst size or normalized frequency, as predicted by the linear constraints. Interestingly, we also observed significant differences in the average expression between the divergent genes group, opposing to the non-divergent group (Fig. S5F).

We then asked if the increased variability in gene expression between species was associated with changes of regulatory complexity (Fig. 5E). Following previous methodology, we selected 28 orthologue genes from the subset of 96 of high coverage genes in mouse and used a genetic algorithm to recapitulate scRNA-seq count distributions with dynamical 2-state and 3-state models (see Materials and Methods and Table S6 for details of the analysis). We then calculated the fold change in the number of conditions (per gene) fitted with 3-state models across all pairwise comparisons of the four species. We found that this fold change correlated (Spearman’s r=0.41, p<0.0001) with the ratio of the slopes between the corresponding linear relationships, such that the transition to a higher slope was associated with increased number of 3-state model fits across corresponding conditions (Fig. 5E). Overall, this demonstrates that evolutionary increases in TLR response variability are associated with increased regulatory complexity, resulting in larger and less frequent transcriptional bursting kinetics.

## Discussion

Transcription is inherently a stochastic process leading to heterogeneity in cell-to-cell mRNA levels. Recent advances suggest the existence of fundamental constraints governing the heterogeneity of gene expression, which rely on the scaling between the variance and mean of the mRNA response distribution [19, 46]. Our previous work, using smFISH data, showed that the overall mRNA variability is linearly constrained by the mean mRNA response across a range of immune-response stimuli [32]. However, these approaches were typically limited by the number of genes considered, not allowing to generalise the observations to the genome-wide scale. Here, utilising an existing scRNA-seq data on the evolutionary-conserved innate immune signalling [4], we perform global analysis of the TLR gene expression response variability and underlying transcriptional bursting. We demonstrate that cell-to-cell variability can be empirically described by a linear function of the population mean across a genome. Based on this, we develop a methodology, relying on statistical modelling of linear mean-variance relationships from single-cell data, that provides a simple yet meaningful way to understand regulation of cellular heterogeneity. We demonstrate that (1) The response heterogeneity of a gene can be defined as the slope of the mean-variance line across >1,500 individual response genes. High variability genes include chemokines and cytokines such as CCL family, while other functional genes are more homogenous, in agreement with previous work [4]. (2) The changes in heterogeneity between species can be described by the change in the slope of the corresponding mean-variance lines, providing insights into the evolutionary control of TLR response variability. (3) The linear relationships determine the underlying transcriptional bursting kinetics, revealing different regulatory modes in response to stimulation and through evolution. (4) Application of dynamical stochastic models of transcription demonstrates a link between the variability and the regulatory complexity, with complexity facilitating heterogeneity via larger and less frequent transcriptional bursting kinetics.

While, in general the available sequencing data are subject to measurement noise [47], and often restricted by the number of data points available, the overall mean-variance relationships were captured using robust linear regression approaches. We first considered regulation of 2,338 TLR-inducible genes in primary murine phagocytes across 20 experimental datasets corresponding to LPS and PIC treatment including biological replicates (Fig. 1). We found that 2,133 relationships were characterised by a significant (non-zero) regression slope (Fig. 1) with 1,551 genes (66% of total) characterised by coefficient of determination *R^2^ >* 0.6. In comparison, out of the 218 genes with one-to-one orthologues between mouse, rat, rabbit and pig, 78% of fitted mean-variance relationships for the 218 genes were characterised by *R^2^ >* 0.6, despite the number of datapoints being limited to 12 (Fig. 5). Fit quality was also reflected in the low mean squared errors between the fitted trends and data, providing good support for the observed phenomenon. We subsequently demonstrated that linear constraints theoretically determine transcriptional bursting characteristics. We used the widely applied moment estimators of the underlying scRNA-seq mRNA distributions to calculate bursting characteristics [1, 3, 11, 18]. Given the empirical linear constraint σ^2^ = *αμ + α_0_*, the burst size and burst frequency become analytical functions of the mean expression (Fig. 2A). We found that 430 relationships (out of 1,551 murine fits) were characterised by statistically significant intercept (*α_0_*). For some genes, this may reflect the empirical nature of these trends, especially for those with small intercept (in relation to variance), for example *Cxcl10* (Fig. 2C). However, we found that many genes with non-zero intercept fits were associated with substantial basal expression in untreated cells, which was also observed previously for the more quantitative smFISH data [32]. Basal expression of the related gene targets has been shown to exhibit different bursting kinetics from the inducible expression [8], which in part may explain the fitted non-zero intercepts for a subset of genes. For *α_0_=0*, linear constraints essentially imply that the burst size must be constant (and equal to the slope of the mean-variance line), while the frequency undergoes modulation with the population mean changes in response to stimulation. This is in general agreement with recent analyses demonstrating a role of frequency in regulation of LPS-induced macrophages [22] or stimulation [9, 20, 48–50]. However, a more detailed investigation of all genes including those with non-zero intercepts, reveals different regulatory modes, including a subset of genes exhibiting burst size modulation (Fig. 3). For instance, a positive intercept is associated with an inverse relationship between the burst size and frequency, while a negative intercept may imply concurrent burst size and frequency changes. As with the mean-variance relationships, the predicted modulation schemes are generally in good agreement with the data in terms of the mean-squared error. Notably, we demonstrate that our methodology can be extended to capture evolutionary differences between species. While gene expression divergence between species has been previously measured in terms of the population response [51], the slope of the linear relationships captures the specific differences in TLR response variability through evolution (Fig. 5). We demonstrate that the evolutionary change of the variability can be described as a ratio *k* between the slopes of the corresponding mean-variance fits, which theoretically implies reciprocal scaling of the burst size and frequency also by *k*. Analysis of the 218 TLR orthologue genes indeed demonstrates that responses of divergent genes are controlled by reciprocal changes of burst size and frequency, while non-divergent genes show the same characteristics across species. Interestingly, we found that within each pair of species, divergent genes exhibited different changes of variability suggesting complex evolutionary traits (e.g., 31 genes exhibiting higher variability in mouse than in pig, and 15 in pig vs. mouse). It would be important to better understand how variability of particular response genes evolved between different species, in the context of their sequence dissimilarities [16, 43–45] as well as epigenetic [52] and signalling components [53] of the TLR signalling between species.

Finally, we used stochastic models of transcription to better understand regulation of transcriptional bursting (Fig. 4). A typical representation involves a 2-state telegraph model, where gene activity changes randomly between “off” and “on” states, facilitating mRNA transcription [1, 3, 18, 36]. However, more complex structures are often used to capture complexity associated with multiple regulatory steps, combinatorial promoter cycling and transcriptional initiation [12, 38, 54, 55]. We hypothesised that TLR response variability is linked with the complexity of the transcriptional regulation. We introduced a 3-state stochastic model, which assumed a sequential activation between “off”, “intermediate” and “on” states, equivalent to promoter cycling [12, 38]. First, we used a computationally efficient Beta-Poisson model, a steady-state approximation of the 2-state telegraph model, which has previously been used to fit scRNA-seq distributions [17, 50]. Values of kinetic parameters were inferred for 7,804 of 31,020 conditions across 1,519 genes demonstrating intermittent transcriptional bursting kinetics [3, 13, 17]. However, this model does not take into account the dynamical nature of the process (measurements at 0, 2, 4 and 6h) and the mRNA half-life with many genes peaking early after stimulation [41]. We therefore used a genetic algorithm to fit the theoretical count distributions to the measured scRNA-seq data using the dynamical 2-state and 3-state models. Based on the Beta-Poisson fits, we selected 96 high coverage murine response genes (and 28 orthologue genes for species analyses), which have existing estimates of mRNA half-life in LPS-stimulated bone marrow derived macrophages [40, 41] or other cell models. These included the highly variable and abundant genes including chemokine family *Ccl5, Ccl4, Ccl3, Ccl2* as well as *IL1b* and *TNFa*. While the scRNA-seq can be in principle treated as time-series data (e.g., across the replicates from individual mice) [34], our current understanding of TLR signalling suggest that due to endotoxin resistance and desensitisation [56–58], the regulatory network, and thus model structures and parameters, are time-varying rather than stationary [59]. We therefore treated each data time-point (and replicate) separately, which also allowed more efficient implementation to fit 1,507 mouse, and 1,079 orthologue conditions. We then used the AIC method [42] to compare the different models considered, and select the one that fitted the measured mRNA distributions most accurately. The results demonstrated that a large subset of genes and conditions fitted a dynamical 3-state model better than the 2-state model. We found that the fraction of conditions explained by the 3-state model correlated well (*r=0.56, p*-value < 0.0001) with slope of the mean-variance relationship, and thus response heterogeneity, for the high coverage murine genes (Fig. 4). Similarly, the increased complexity was associated with evolutionary changes of response variability between species (Fig. 5). In general, we found that increased regulatory complexity facilitated larger response variability through increased burst sizes and reduced frequency of transcriptional bursting (Fig. 4D), while scRNA-seq count variance exhibited correlations with transcription rates and ‘on’ rates. A better understanding of the relationships, and in particular mechanistic basics for controlling gene-specific slopes (i.e., response variability) as well as their sensitivity to pharmacological perturbation and infection and disease state, will require further detailed investigations [22]. Nevertheless, we believe that our methodology, relying on the inference of mean-variance relationships, provides new insight into regulation of single-cell variability of innate immune signalling and will be applicable to other inducible gene expression systems.

## Materials and Methods

### Analysis environment

Computational analysis was performed using Python v3.8.2 in a 64-bit Ubuntu environment running under Windows Subsystem for Linux (WSL) 2 and using the conda v4.8.3 package manager. Relevant packages were NumPy v1.19.1 (Van Der Walt *et al*., 2011), pandas v1.0.5 (Reback *et al*., 2020), Scanpy v1.5.1 (Wolf *et al*., 2018), scikit-learn v0.23.1 (Pedregosa *et al*., 2011), SciPy v1.4.1 (Virtanen *et al*., 2020) and statsmodels v0.11.1 (Seabold and Perktold, 2010) for processing and Matplotlib v3.2.1 (Hunter, 2007) and seaborn v0.10.1 (Waskom *et al*., 2020) for visualisation. Robust linear regression models and Benjamini-Hochberg false discovery rate (FDR) correction was performed in statsmodels. Coefficient of determination (*R*^2^) scores were calculated using the metrics module of scikit-learn.

### Acquisition and processing of mRNA count data

mRNA count data associated with the study by Hagai *et al*. (2018) were downloaded from the Array Express database, in particular, the E-MTAB-6754.processed.2.zip file to obtain the UMI counts of bone marrow-derived mononuclear phagocytes from mouse, rat, pig and rabbit. Phagocytes were either untreated (0h) or stimulated with LPS for 2, 4 and 6 h, resulting in 12 scRNA-seq datasets per species. In addition, phagocytes from mice and rat were also treated with PIC at 2, 4 and 6h. Notably, the dataset contains no UMI counts for PIC stimulation at 6 h for mouse 1 but has two for mouse 2 (labelled 6 and 6A). When collating the counts, the missing replicate for mouse 1 was disregarded and the PIC 6A time point – assumed to be a technical replicate – was excluded. Therefore, 20 datasets (referred as conditions herein) for the mouse, 21 datasets for the rat, 12 conditions for the pig and the rabbit dataset were considered for each gene (see Table S4). The UMI counts were median scaled per cell using the normalize_total function of Scanpy and subsequently used for fitting mean-variance relationships and bursting modulation. Integer values, referred to as “mRNA counts” in this work were used for mathematical model fitting (see Github repository for data normalisation, UMI normalisation [60] and extraction of mRNA count distributions). Gene IDs, gene symbols and the descriptions of the genes were obtained from the Ensembl Release 103 database of the four studied species: *Mus musculus* (mouse), *Rattus norvegicus* (rat), Sus scrofa (pig) and Oryctolagus Cuniculus (rabbit) using the BioMart web tool (Yates *et al*. 2020). Hagai *et al*. (2018) defined a set of 2,336 LPS-responsive genes based on differential expression in response to LPS stimulation with FDR-corrected *p*-value < 0.01 and existing orthologues in rabbit, rat and pig. *Il1b* and *Tnf* were added to this list – as well characterised TLR-response genes from the study of Bagnall *et al*. (2020)–resulting in a set of 2,338 LPS response genes with 46,740 conditions overall. Similarly, the responsive genes from the three other species were also determined. 2586 rat genes, 1892 pig genes and 859 rabbit genes showed differential expression upon LPS stimulus. 218 one-to-one orthologue genes were found to be responsive in all species, these genes formed the analysis subset.

### Fitting theoretical bursting characteristics

The sample mean (μ) and variance (σ^2^) of mRNA counts were calculated for the measured mRNA count distribution for individual response genes across conditions. The mean-variance relationships (σ^2^ = α*μ + α*_0_) were fitted using robust linear regression, using a Huber M-estimator with a tuning constant of 1.345, across all relevant conditions. A model’s fit was considered successful if the slope (*α*) was statistically significant based on FDR-adjusted *p*-value < 0.05, and it provided a good overall fit (unweighted *R^2^* > 0.6). FDR-adjusted *p*-value < 0.05 was also calculated for the intercept (*α*_0_). Assuming linear constraints of mRNA mean and variance, theoretical bursting characteristics were analytically derived, using moment estimators; burst size *b_s_=α_0_/μ + α*, burst frequency *b_f_=μ^2^/(α_0_+αμ)* and *b_f_=α0/(b_s_(b_s_-α))*. Relative root mean square error, 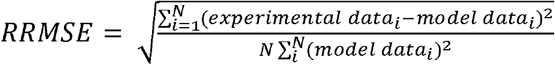, where *N* denoted the number of datapoints, was used to compare theoretical predictions and experimental data. Relative fold change was used to calculate the level of burst size and frequency modulation in the measured data, across all the conditions per gene:

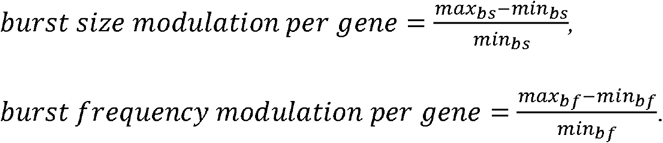

Comparison between burst size and burst frequency modulation was quantified as the ratio of the two quantities, i.e., 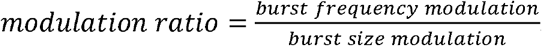.

### Pairwise comparison of the slopes of the mean-variance regressions

The differences in the mean-variance relationships of a gene between species were measured by pairwise comparisons between the slopes. A Student’s t-test was performed to determine whether the two slopes are statistically significantly different, or not. The following formula was used to calculate the t-statistic values:

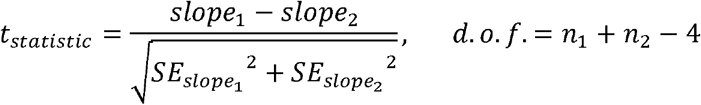

SE_slope_ represents the standard error of the value of the slope in the fitting of the robust linear regression model on the data. The degrees of freedom (d.o.f.) is dependent on the number of data points used to create the two linear regression lines compared (*n_1_* and *n_2_*, respectively). *p*-values were determined using the cumulative distribution function of the relevant t distribution. As the four slopes were compared pairwise, six *p*-values were calculated per gene. *p*-values were corrected by the Benjamini-Hochberg procedure. Two slopes were deemed significantly different if the false discovery rate (FDR) corrected *p*-value was below 0.05. Subset of genes with different number of significant FDR-corrected *p*-values were compared using a measure of evolutionary response divergence, such that *response divergence = log[1/3 × ∑_j_(log[FC pig] - log[FC glire_j_])^2^]*, with j =(1,2,3) corresponding to 3 glires (mouse, rat and rabbit) and FC is the fold change in response to LPS stimulation per gene (Supplementary Table 4 in [4]).

### Inference of Beta-Poisson model

Inference of Beta-Poisson model parameters (*k_on_, k_off_* and *k_t_*) from individual scRNA-seq count distributions was performed using the profile-likelihood txburstML script (Larsson *et al*., 2019) downloaded from GitHub (version 1844c47be5f1ad2104cf15d425889768ec45df8b). Conditions that txburstML did not mark as “keep” (indicating convergence) were discarded. Genes with a least 10 fitted conditions per mouse (out of 20) and rat (out of 21) as well at least 6 in the pig and rabbit (out of 12) were included in the high coverage gene sets.

### Modelling and inference of dynamical models of transcription

Theoretical temporal mRNA distributions for considered models of transcription were obtained using the Chemical Master Equation (CME) following our previous approach [32]. In brief, the time evolution of the probability distribution over mRNA counts *P*(***X**,t*), is given by *P*(***X**, t*) = exp[*R*(*Θ*)*t*] *P*_0_(**X**), where *R*(*θ*) is a transition rate matrix describing flow of probability between different states, where a state is defined by the number of mRNA in the cell at time *t* and the transcriptional states of the gene’s alleles. *P*_0_(*X*) is specified by initial data such that ∑_*x*_*P*_0_(***X***) = 1. *P*(***X***, *t*) is calculated using a fast matrix exponential function implemented in MATLAB by [61]. All simulations begin with initial conditions of no mRNA and both gene alleles being in the *‘off* state. *R*(*θ*) depends on model structure and the parameters. In this work, we considered a *stochastic telegraph model*—with two independent alleles per gene, the activity of which switches randomly between *‘off* and ‘*on’* states, with the latter being permissive for mRNA transcription [1, 3, 36, 62]. The associated kinetic parameters include switching ‘*on*’ and *“off* rates (*k_on_* and *k_off_*, respectively) as well as rates of mRNA transcription and degradation (*k_t_* and *k_d_*, respectively). We also considered an extended model including an additional regulatory step, such that each allele exists in one of three states: an inactive *‘off*, an intermediate ‘*I’* or an active ‘*on*’. Reversible stochastic transitions (with appropriate rates) occur between the inactive and intermediate (*t_on_* and *t_off_*), the intermediate and active states (*k_on_* and *k_off_*), as well as direct transition between active and inactive states (*k_c_*). We further assume that transcription occurs only in the intermediate and active states (*k_ti_* and *k_t_*, respectively).

A genetic algorithm (GA) was implemented using the *ga* function in MATLAB and employed to estimate model parameters. We minimized an objective function given by the average absolute distance between the theoretical (CME) and measured cumulative distribution functions (CDFs) across observed mRNA counts per condition 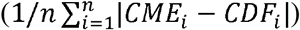, where *i’s* are unique mRNA counts observed in the measured distributions (for those with total unique counts n>1). CDFs were calculated using empirical cumulative distribution function (*ecdf*). The best of 10 model fits from independent GA runs for each condition (using a population size of 100, elite count of 2, crossover factor of 0.6, 20 generations and the tournament selection function) was retained. Gene activation/inactivation rates were constrained between 0 and 1 min^-1^, transcription was constrained between 0 and 50 mRNA counts min^-1^ per allele, which is the same order of magnitude to previous estimates [2, 3, 62, 63]. Murine mRNA half-lives were obtained from literature, when available from LPS-stimulated bone marrow derived macrophages [40, 41] or other cell models [64–71]. Murine half-lives were also used when fitting orthologue genes.

Akaike’s Information Criterium (AIC) was used to asses model fits and perform model selection [42]. *AIC = 2p* — 21og[*L*(Θ|*X*)] where log [*L*(Θ|*X*)] is the log-likelihood function of the fitted mRNA count distribution given measured data *X* defined as 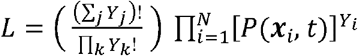 with *Y^t^* being a vector of the number of cells displaying each observed state at time t (the sum of this vector is the total number of cells N), and *p* corresponds to number of parameters in the model; resulting in a penalty for higher complexity. Models with AIC larger than Q3+1.5(Q3-Q1), where Q1 and Q3 are the first and third quartiles of the AIC distribution per model across genes were removed. As a result, out of 1507 mouse, and 1079 orthologue (pig, rat and rabbit) conditions, 1210 and 981 that fitted 2- and 3-state models were retained, respectively.

## Statistical analyses

Statistical analysis was performed using GraphPad Prism 8 software (version 8.4.2). The D’Agostino-Pearson test was applied to test for normal (Gaussian) distribution of acquired data. Two-sample comparison was conducted using non-parametric Mann Whitney test. For analyses of variance Kruskal-Wallis ANOVA with Dunn’s multiple comparisons test was performed. Coefficient of determination (*R^2^*) was used to assess regression fits; Spearman correlation coefficient *r* was used to test association between other variables.

## Author contributions

NA performed analyses presented in the manuscript. DN and ZW performed preliminary analyses and developed Python codes. MM and PP provided supervision and conceptualisation. PP with assistance of NA and MM wrote the manuscript. All authors read and approved the final manuscript.

## Data Availability Statement

Python codes developed in this study are available from Github repository (https://github.com/ppaszek/TLR_bursting).

---

**Figure S1. Analysis of the variability in the TLR responses. A.** Fitted regression lines for the 1,551 high confidence genes, shown are genes with different range of the slope α. Highlighted in different colours are fits for the individual genes. Broken line indicates μ=σ^2^ line. **B**. Histogram of the measured mRNA response range for the 1,551 high confidence genes. **C**. Effect of the slope (left) and intercept (right) of the mean-variance relationship on the burst size and burst frequency modulation. Shown are simulated burst size and frequency modulation schemes for a range of α and α_0_ (as indicated on the graph). **D**. Modulation schemes for *Jchain* gene. Shown is the comparison between theoretical relationships based on fitted mean-variance relationships (in red) and corresponding estimates from data (open circles). Equation for fitted mean-variance relationships highlighted in the top left panel, respectively. **E**. Relationship between the slope (α) and in the intercept (α_0_) across fitted 1,551 high confidence genes.

**Figure S2. Inferred kinetic parameter rates for 2-state telegraph model using Beta-Poisson model. A.** Comparison between the 1,551 high confidence genes across all conditions that either fit or do not fit the Beta-Poisson model. **B.** Histogram of fitted *k_on_, k_off_* and *k_t_* across 7704 conditions for 1,519 high confidence genes. Inference performed using profile likelihood of the Beta-Poisson model. Parameters units are expressed per degradation half-life **C**. Relationship between inferred *k_on_* vs. *k_off_* rates (left) and *k_on_* vs. *k_t_* (right) across parameters from A. Rates for *Nfkbia, Il12* and *Ccl5* highlighted in different colours. Identity line depicted with a broken line. **D.** Histogram of the number of inferred conditions across 1,159 high confidence genes. Broken line highlights the threshold for at least 10 conditions fitted per gene. **E.** Histogram of the fitted regression slopes for the 96 high coverage gene set. **F.** Fitted regression lines for the 96 high coverage genes. Highlighted in colour are fits for the individual genes of interest. Broken line indicates *μ=σ^2^* line.

**Figure S3. Analysis of stochastic models of transcription. A**. Comparison between the fitted and measured scRNA-seq count distributions for few gene examples. Shown are cumulative probability distribution of data (in green) vs. the corresponding Beta-Poisson, 2-state and 3-state model fits (in blue, red and violet, respectively) for *Adm* (LPS, 2h, replicate 1), *Il1α* (PIC, 2h, replicate 1), *Cd40* (LPS, 4h, replicate 1) and *Il7r* (0h, replicate 2) genes. Ratios of respective AICs between models highlighted on top. **B**. Summary of comparing Beta-Poisson, 2-state and 3-state model fits across the conditions of the high coverage genes. Best models defined either by AIC smaller (in white) or 2-fold smaller (in black) than the next best model. **C**. Summary of 2-state and 3-state model fits across a range of thresholds *T= AIC_2-state_/AIC_3-state_* for the fitted 96 high coverage genes across all conditions. **D.** Relationships between the number of Beta-Poisson, 2-state and 3-state model fits for the 96 high coverage genes across all conditions. Best fit model defined by *AIC_best model_<AIC_2nd best_*.

**Figure S4. Model-based analysis of transcriptional bursting. A**. Summary of 2-state model fits defined for 141 conditions such that *AIC_2.state_<0.5AIC_3.state_* (as in Fig. 4C). Shown is the distribution of fitted *k_on_* (min^-1^) and *k_off_* (min^-1^) rates as well as Spearman correlation coefficient r with mRNA variance. **B**. Summary of 3-state model fits defined for 266 conditions such that *AIC_3-state_<0.5AIC_2-state_* (as in Fig. 4C). Shown is the distribution of fitted rates as well as Spearman correlation coefficient r with mRNA variance (and between selected rates). **C**. Comparison between fitted transcription rates for 2-state and 3-state models (as in A and B, respectively). Statistical significance assessed with Kruskall-Wallis test with Dunn’s correction for multiple comparisons (* *p*-value < 0.05, *** *p*-value < 0.001). **D**. Analysis of transcriptional bursting across high coverage genes and conditions fitted by 2-state vs 3-state models. Shown is the comparison between best fit 2- and 3-state models in terms of mean mRNA expression, variance, burst size and frequency. Best fit defined by *AIC_best model_<AIC_2nd best_* (from Fig. S3B). Burst size and frequency calculated per condition using moment estimators. Statistical significance assessed with Mann-Whitney test (* *p*-value < 0.05, *** *p*-value < 0.001, **** *p*-value <0.0001, ns not significant).

**Figure S5. Analysis of transcriptional bursting across species. A.** Schematic diagram of data analysis; 169 orthologue genes exhibiting good mean-variance fits (*R^2^ >* 0.6) statistically tested for differences in the slope of the linear fit. Right: Venn diagram of TLR response orthologue genes in at least one of the species studied by Hagai et al. (2018). **B**. Fitted mean-variance relationships for a subset of orthologue genes across species. Shown is the comparison between the fitted mean-variance relationships (in solid lines, colour-coded by species) and corresponding data (circles). **C**. Evolutionary response divergence across orthologue gene subsets defined by the number of statistically significant FDRs between fitted regression slopes across four species (as in Table S2). Statistical significance assessed using ordinary ANOVA with Dunnett’s correction for multiple comparisons (*** *p*-value < 0.001, * *p*-value < 0.05, ns not significant). **D**. Global modulation of transcriptional busting across species. Shown is the comparison between fitted mean-variance relationship and theoretical burst size and frequency modulation schemes vs. relationships derived from data. Shown is a violin plot of relative root mean square error (RRMSE) of 169 orthologue genes. **E**. Histogram of the slope ratio (α_mouse_/α_pig_) for the 169 orthologue genes between mouse and pig. α_mouse_ and α_pig_ denote slopes of the fitted mean-variance relationships for each pair of species per gene. **F**. Analysis of divergent and non-divergent mouse and pig TLR-response genes. Shown are box plots of average mRNA expression per gene stratified into divergent (α_mouse_> 2α_pig_ or α_pig_>2α_pig_) and complementary non-divergent group (31, 15 and 123 orthologue genes, respectively). Statistical significance assessed with a paired Wilcoxon test (*** *p*-value < 0.001,ns not significant).

**Table S1.** Fitted mean-variance relationships for the mouse TRL response genes.

**Table S2.** Modulation of transcriptional bursting across 1,551 mouse high confidence genes. **Table S3.** Modelling of scRNA-seq count distributions.

**Table S4.** Number of phagocyte cells and genes measured in each single cell in the four species. Only the genes showing expression under at least one condition were studied **Table S5.** Pairwise comparison of the slopes of the mean-variance regression lines was performed between each two species. The table shows the number of significant FDR values (<0.05) obtained for each of the 169 orthologue genes studied.

**Table S6.** Analysis of TLR response variability across species.

