## Supplementary Figures for "Variability of the innate immune response is globally constrained by transcriptional bursting"

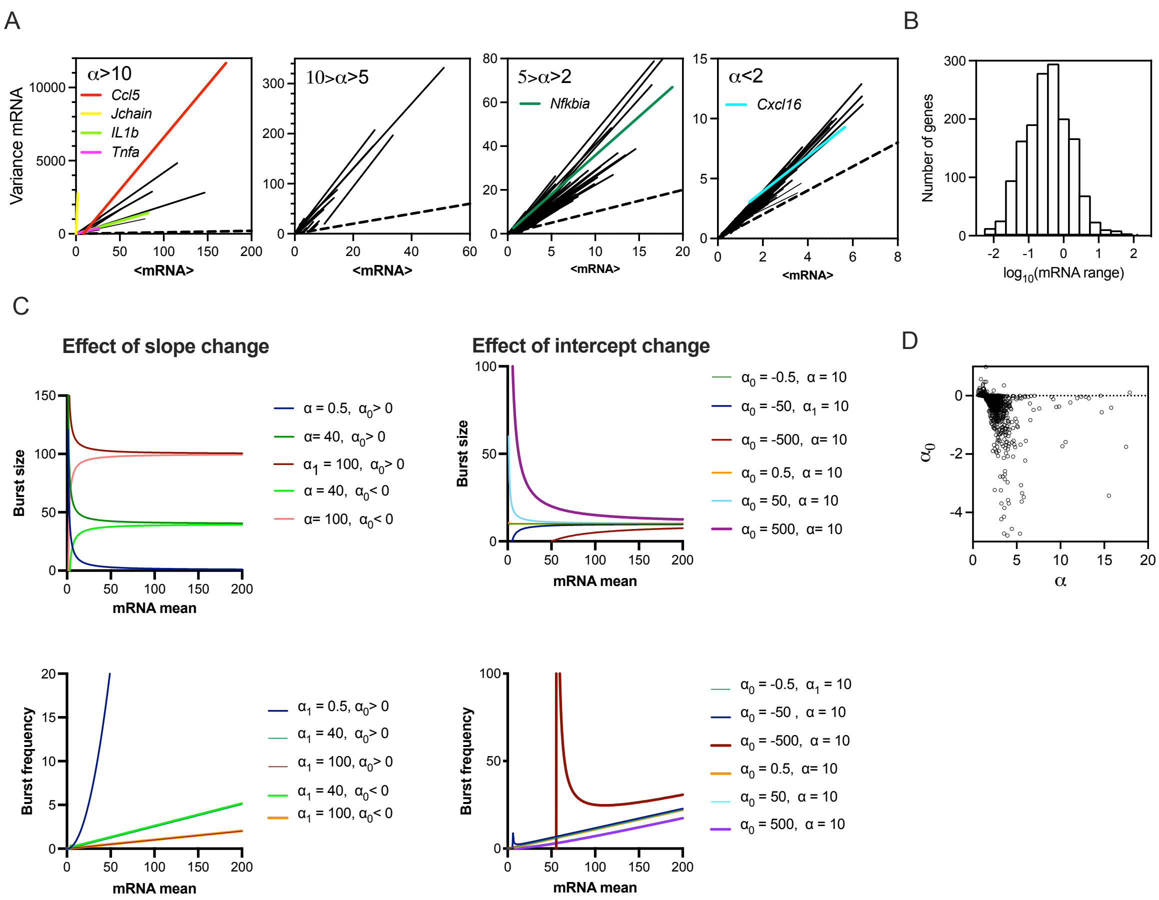


**Figure S1. Analysis of the variability in the TLR responses. A.** Fitted regression lines for the 1,551 high confidence genes, shown are genes with different range of the slope α. Highlighted in different colours are fits for the individual genes. Broken line indicates $\mu=\sigma^2$ line. **B**. Histogram of the measured mRNA response range for the 1,551 high confidence genes. **C**. Effect of the slope (left) and intercept (right) of the mean-variance relationship on the burst size and burst frequency modulation. Shown are simulated burst size and frequency modulation schemes for a range of α and $\alpha_0$ (as indicated on the graph). **D**. Modulation schemes for *Jchain* gene. Shown is the comparison between theoretical relationships based of fitted mean-variance relationships (in red) and corresponding estimates from data (open circles). Equation for fitted mean-variance relationships highlighted in the top left panel, respectively. **E**. Relationship between the slope (α) and in the intercept ($\alpha_0$) across fitted 1,551 high confidence genes.

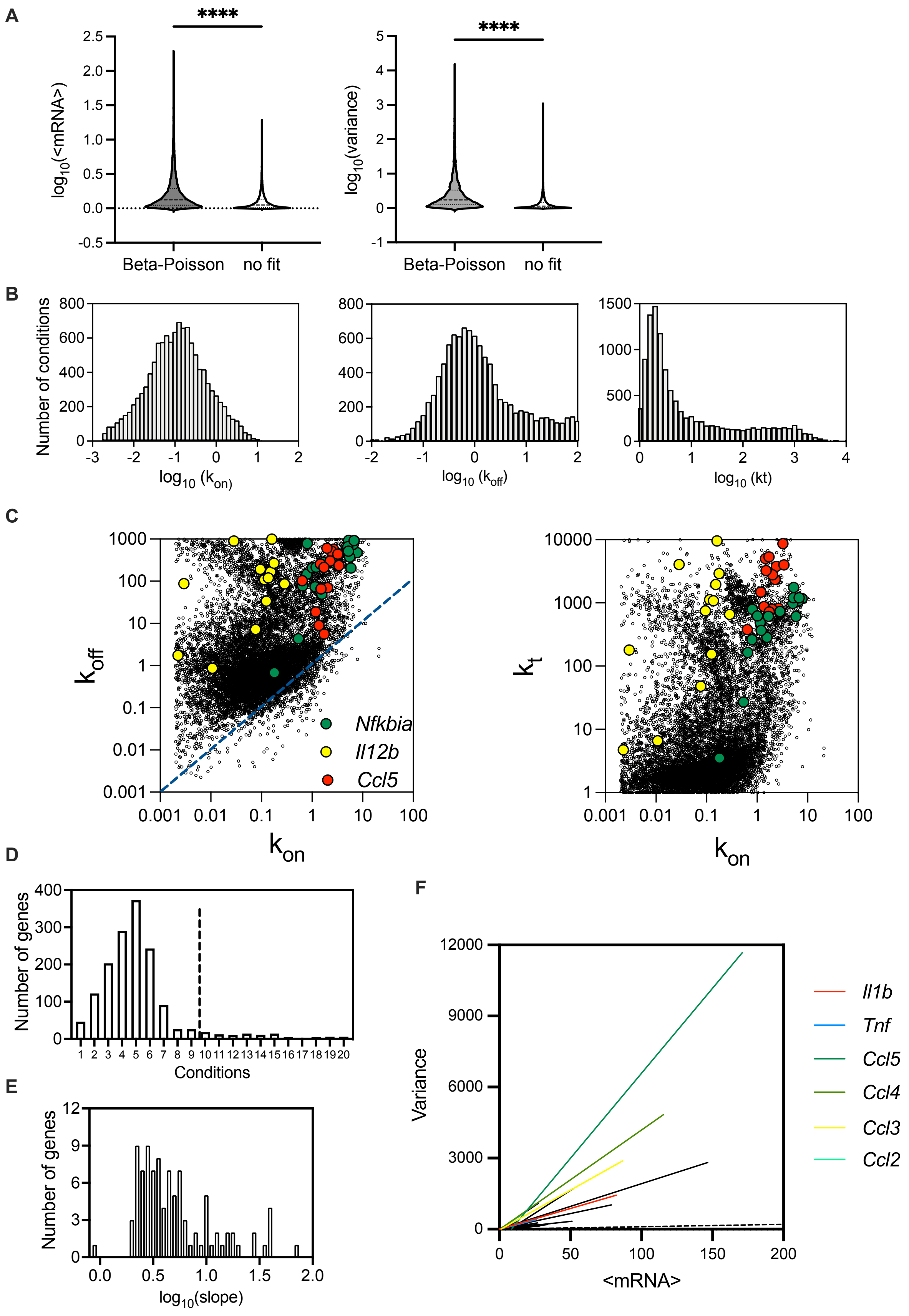

A
****
log10(<mRNA>)
Beta-Poisson
no fit
log10(variance)
B
Number of conditions
log10 (kon)
log10 (koff)
log10 (kt)
C
koff
kon
Nfkbia
Il12b
Ccl5
kt
D
Number of genes
Conditions
E
Number of genes
log10(slope)
F
Variance
<mRNA>
Il1b
Tnf
Ccl5
Ccl4
Ccl3
Ccl2

**Figure S2. Inferred kinetic parameter rates for two-state telegraph model using Beta-Poisson model. A.** Comparison between the 1551 high confidence genes across all conditions that either fit or do not fit the Beta-Poisson model. **B.** Histogram of fitted $k_{on}$, $k_{off}$ and $k_t$ across 7704 conditions for 1,519 high confidence genes. Inference performed using profile likelihood of the Beta-Poisson model. Parameters units are expressed per degradation half-life **C**. Relationship between inferred $k_{on}$ vs. $k_{off}$ rates (left) and $k_{on}$ vs. $k_t$ (right) across parameters from A. Rates for *Il12*, *Nfkbia* and *Ccl5* highlighted in different colours. Identity line depicted with a broken line. **D.** Histogram of the number of inferred conditions across 1,159 high confidence genes. Broken line highlights the threshold for at least 10 conditions fitted per gene. **E.** Histogram of the fitted regression slopes for the 96 high coverage gene set. **F.** Fitted regression lines for the 96 high coverage genes. Highlighted in colour are fits for the individual genes of interest. Broken line indicates $\mu=\sigma^2$ line.

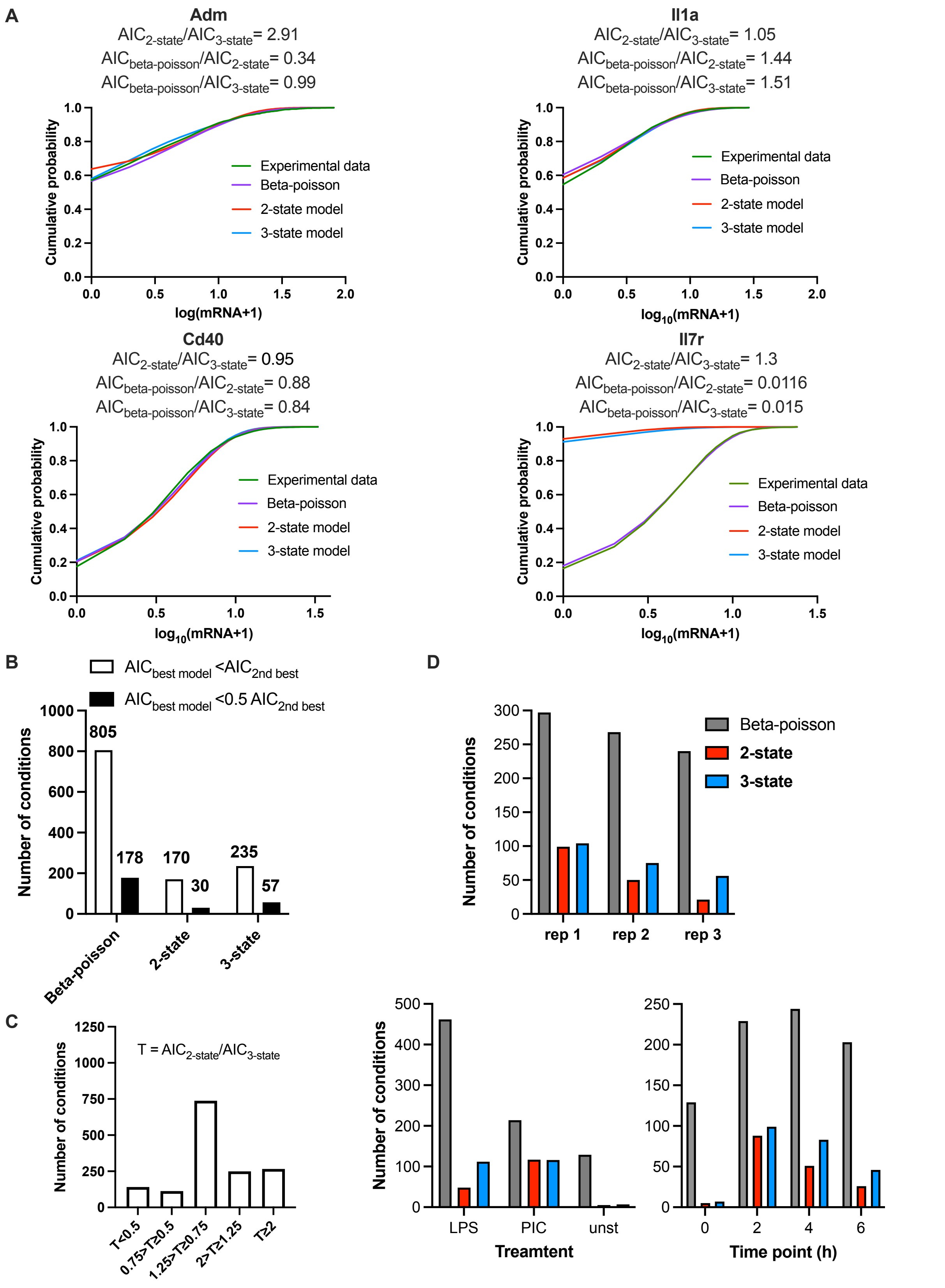


**Figure S3. Analysis of stochastic models of transcription. A**. Comparison between the fitted and measured scRNA-seq count distributions for few gene examples. Shown are cumulative probability distribution of data (in green) vs. the corresponding Beta-Poisson, 2-state and 3-

state model fits (in blue, red and violet, respectively) for *Adm* (LPS, 2h, replicate 1), *Il1α* (PIC, 2h, replicate 1), *Cd40* (LPS, 4h, replicate 1) and *Il7r* (0h, replicate 2) genes. Ratios of respective AICs between models highlighted on top. **B**. Summary of comparing Beta-Poisson, 2-state and 3-state model fits across the high coverage genes and conditions. Best models defined either by AIC smaller (in white) or 2-fold smaller (in black) than the next best model. **C**. Summary of 2- and 3-state model fits across a range of thresholds $T = AIC_{2\text{-}state}/AIC_{3\text{-}state}$ for the fitted 96 high coverage genes across all conditions. **D.** Relationships between the number of Beta-Poisson, 2-state and 3-state model fits for the 96 high coverage genes across all conditions. Best fit model defined by $AIC_{best\ model} < AIC_{2nd\ best}$.

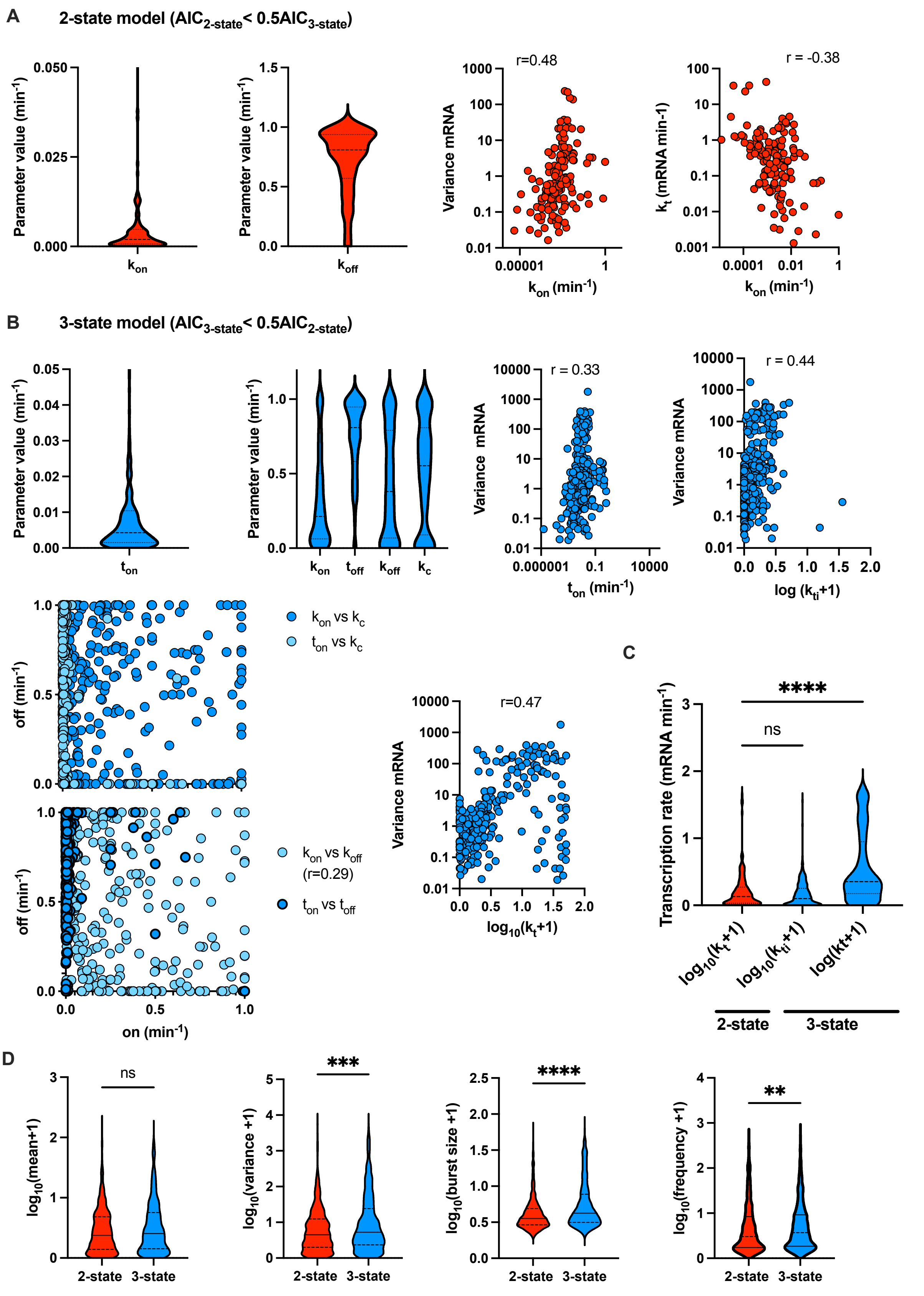


**Figure S4. Model-based analysis of transcriptional bursting.** **A**. Summary of 2-state model fits defined for 173 conditions such that $AIC_{2\text{-}state}<0.5AIC_{3\text{-}state}$ (as in Fig. 4C). Shown is the distribution of fitted $k_{on}$ (min$^{-1}$) and $k_{off}$ (min$^{-1}$) rates as well as Spearman correlation coefficient r with mRNA variance. **B**. Summary of 3-state model fits defined for 275 conditions such that $AIC_{3\text{-}state}<0.5AIC_{2\text{-}state}$ (as in Fig. 4B). Shown is the distribution of fitted rates as well as Spearman correlation coefficient r with mRNA variance (and between selected rates). **C**. Comparison between fitted transcription rates for 2-state and 3-state models (as in A and B, respectively). Statistical significance assessed with Kruskall-Wallis test with Dunn's correction for multiple comparisons (* p value<0.05, *** p value <0.001). **D**. Analysis of transcriptional bursting across high coverage genes and conditions fitted by 2-state vs 3-state models. Shown is the comparison between best fit 2- and 3-state models in terms of mean mRNA expression, variance, burst size and frequency. Best fit defined by $AIC_{best\ model}<AIC_{2nd\ best}$ (from Fig. S3C). Burst size and frequency calculated per condition using moment estimators. Statistical significance assessed with Mann-Whitney test (* p value<0.05, **** p value <0.0001, ns- not significant).

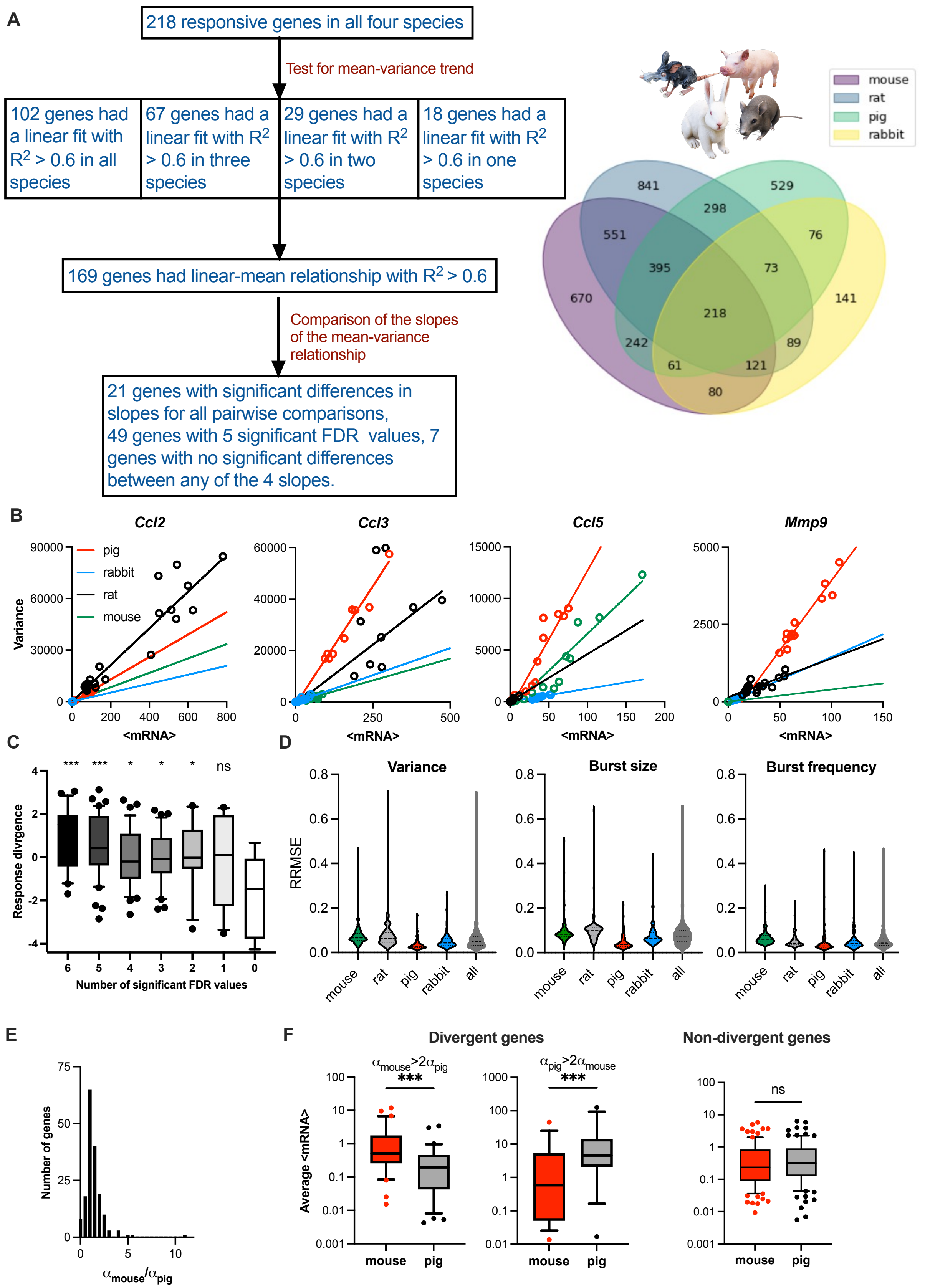


**Figure S5. Analysis of transcriptional bursting across species. A.** Schematic diagram of data analysis; 169 orthologue genes exhibiting good mean-variance fits ($R^2>0.6$) statistically tested for differences in the slope of the linear fit. Right: Venn diagram of TLR response orthologue genes in at least one of the species studied by Hagai et al. (2018). **B**. Fitted mean-variance relationships for a subset of orthologue genes across species. Shown is the comparison between the fitted mean-variance relationships (in solid lines, colour-coded by species) and corresponding data (circles). **C**. Evolutionary response divergence across orthologue gene subsets defined by the number of statistically significant FDRs between fitted regression slopes across four species (as in Table S2). Statistical significance assessed using ordinary ANOVA with Dunnett's correction for multiple comparisons (*** p-val<0.001, * pval<0.05, ns – not significant). **D**. Global modulation of transcriptional busting across species. Shown is the comparison between fitted mean-variance relationship and theoretical burst size and frequency modulation schemes vs. relationships derived from data. Shown is a violin plot of relative root mean square error (RRMSE) of 169 orthologue genes. **E**. Histogram of the slope ratio ($\alpha_{mouse}/\alpha_{pig}$) for the 169 orthologue genes between mouse and pig. $\alpha_{mouse}$ and $\alpha_{pig}$ denote slopes of the fitted mean-variance relationships for each pair of species per gene. **F**. Analysis of divergent and non-divergent mouse and pig TLR-response genes. Shown are box plots of average mRNA expression per gene stratified into divergent ($\alpha_{mouse}> 2\alpha_{pig}$ or $\alpha_{pig}>2\alpha_{pig}$) and complementary non-divergent group (31, 15 and 123 orthologue genes, respectively). Statistical significance assessed with a paired Wilcoxon test (**** p-val<0.0001, *** p-val <0.001, ns not significant).
