## Supplementary Table 4 for "Variability of the innate immune response is globally constrained by transcriptional bursting"

| Species | Number of cells | Total number of genes | Genes showing expression | Number of conditions |
|---|---|---|---|---|
| Mouse | 53086 | 22048 | 16798 | 20 |
| Rat | 50185 | 22277 | 16780 | 21 |
| Pig | 23469 | 21607 | 15602 | 12 |
| Rabbit | 34528 | 19293 | 14480 | 12 |

**Table S4.** Number of phagocyte cells and genes measured in each single cell in the four species. Only the genes showing expression under at least one condition were studied
