## Supplementary Table 5 for "Variability of the innate immune response is globally constrained by transcriptional bursting"

| Number of significant FDR values | Genes |
|---|---|
| 6 | *Car4, Ccl2, Ccl4, Ccl5, Cxcl10, Ehd1, F3, Ier3, Ifit2, Ifnb1, Inhba, N4bp1, Nampt, Nlrp3, Parp9, Sema3c, Serpinb2, Slamf7, Tagln2, Tnfaip3, Tnfsf15* |
| 5 | *Adora2a, Arrdc3, Atad1, Cblb, Ccl20, Ccl3, Ccrl2, Cflar, Cmpk2, Csrnp1, Fam105a, Hmgcs1, Ifi44, Il10, Il1a, Il27, Il4ra, Irf1, Mef2c, Mmp3, Mmp9, Mxd1, Nabp1, Nfkbiz, Nrp2, Nub1, Parp11, Pik3ap1, Pim1, Rab32, Rasgef1b, Rel, Rnd1, Rnf19a, Sdc4, Serpinb8, Slamf1, Slc46a3, Snx10, Socs1, Srgn, Stat3, Tcf7l2, Tfec, Tnfaip6, Tnfsf10, Ttc39b, Txnip, Zc3hav1* |
| 4 | *A230050P20Rik, Acsl1, Cd274, Cd40, Cd53, Cdkn2c, Cxcl9, Cxcr4, Dusp2, Fas, Fgd4, Fmr1, Lpxn, Manf, Marcks, Mov10, Nbr1, Nr3c1, Olr1, Plekho1, Ppa1, Ppp1r15a, Psma6, Rgs10, Samsn1, Slc17a5, Slc29a3, Slc37a2, Tiparp, Tnip1, Tra2a, Trim25, Ulk1, Vcan, Ypel3* |
| 3 | *Amacr, Arl5b, Atp10a, Birc3, Ccng2, Coprs, Gmnn, Hbegf, Hhex, Icam1, Jak2, Mafb, Mb21d1, Mical1, Mxd4, Nfkbia, Nmi, Npc1, Nr1d2, Nr4a3, Pcgf5, Plk2, Pnrc1, Rnd3, Rnf19b, Sh3pxd2b, Smarca2, Tdrd7, Tfdp2, Traf3ip2, Trim26, Uap1, Wars, Xrn1* |
| 2 | *Baz1a, Ccdc34, Gmpr, Nfkb2, Nfkbib, Plekhm1, Prkag2, Rybp, Skil, Tmem51, Uqcc3, Xpc* |
| 1 | *Csrp2, Fam98c, Frmd4b, Gtf2i, Ldlrap1, Lpar6, Mapk6, Rasa2, Rragd, St6gal1, Top1* |
| 0 | *Arhgap4, Camk2g, Cbx8, Crot, Hdac5, Tmbim6, Uri1* |

**Table S5.** Pairwise comparison of the slopes of the mean-variance regression lines was performed between each two species. The table shows the number of significant FDR values (<0.05) obtained for each of the 169 orthologue genes studied.
